# CAKUT variants in *PRPF8, DYRK2*, and *CEP78*: implications for splicing and ciliogenesis

**DOI:** 10.1101/2025.07.16.665151

**Authors:** Lea M Merz, Shirlee Shril, Tucker J. Carrocci, Csenge K. Rezi, Natalie J. Zeps, Rafael Jiménez-Izquierdo, Florian Bergmann, Narcis Adrian Petriman, Caroline M Kolvenbach, Nils D Mertens, Søren L. Johansen, Jan Halbritter, Alina Christine Hilger, Shaikh Qureshi Wasay Mohiuddin, Kathryn E Hentges, Hila Milo Rasouly, Ali G Gharavi, Kiyotsugu Yoshida, Esben Lorentzen, Marco Calzado, Andreas Kispert, Saishu Yoshida, Lotte B. Pedersen, Aaron A. Hoskins, Florian Buerger, Friedhelm Hildebrandt

## Abstract

**Introduction:** Congenital anomalies of the kidney and urinary tract (CAKUT) are the leading cause of chronic kidney disease in children and young adults. Although over 50 monogenic causes have been identified, many remain unresolved. PRPF8 is a core spliceosome component, essential for pre-mRNA splicing, and further localizes to the distal mother centriole to promote ciliogenesis.

**Methods:** We performed trio exome sequencing in 208 CAKUT families and identified strong variants in *PRPF8* and the EDD-DYRK2-DDB1^VprBP^ complex. Functional validation included splicing assays in yeast (Saccharomyces cerevisiae), Sonic hedgehog (Shh) signaling in RPE-1 cells, co-immunoprecipitation for protein complex assembly, and in situ hybridization in mouse embryos. Protein interactions were modeled using AlphaFold.

**Results:** We identified heterozygous *de novo* or inherited variants in *PRPF8, DYRK2, DDB1, EDD* and *CEP78.* Yeast assays revealed that while most PRPF8 variants preserved growth and splicing at consensus splice sites, the de novo *PRPF8*^R1681W^ variant impaired splicing of non-consensus splice sites and was inviable at elevated temperature. CAKUT variants failed to rescue *prp28-1* and *U4-cs1* alleles but showed variant-specific synthetic interactions with *brr2-1*, including weak suppression or synthetic sickness at elevated temperatures. Shh signaling was reduced in ∼50% of *PRPF8* variants expressed in RPE-1 cells. *CEP78* truncating variants abrogated binding to CEP350 and VPRBP. Two *DYRK2* variants disrupted EDD-DYRK2-DDB1VprBP complex formation without affecting kinase activity. In situ hybridization revealed strong *Prpf8* expression in the developing collecting duct and urothelium.

**Conclusion:** Variants in *PRPF8* and components of the EDD-DYRK2-DDB1^VprBP^ complex may contribute to CAKUT through impaired pre-mRNA splicing and defective ciliogenesis. These findings uncover an entirely new functional network of candidate genes for CAKUT and ciliopathies, significantly broadening our understanding of disease mechanisms and offering novel entry points for mechanistic studies.

**Translational Statement:** Our study identifies a previously unrecognized molecular network involving PRPF8 and the EDD–DYRK2–DDB1VprBP complex, revealing a novel pathogenic mechanism in CAKUT. These results introduce a new class of candidate genes and pathways essential for kidney development. As the genetic etiology of CAKUT remains unknown in most patients, our findings underscore the need for targeted genetic testing and functional studies to enhance diagnosis, advance mechanistic insight, and enable more personalized clinical management.

## INTRODUCTION

Congenital anomalies of the kidney and urinary tract (CAKUT) are common birth defects and the leading cause (∼40%) of chronic kidney disease in individuals under 30 years of age^1^. Around 50 monogenic CAKUT genes have been identified, enhancing our understanding of kidney development. Ciliopathies, caused by defects in cilia-related genes, share overlapping kidney phenotypes with CAKUT, including hydronephrosis and hypoplastic kidneys, often with extra-renal features and broad phenotypic variability^6^.

PRPF8 (Pre-mRNA-processing-splicing factor 8) is a highly conserved core component of the U5 snRNP subcomplex of the spliceosome **(Fig. 1A)**^2–4^. It forms a protein scaffold supporting the U2/U6 snRNA catalytic center and drives conformational rearrangements necessary for pre-mRNA splicing **(Fig. 1A)**^7–10^. *PRPF8* variants can disrupt gene expression programs linked to cell growth, mitosis, and the cell cycle^11,12^. Its molecular function and interactions with splicing factors like Brr2, U4 snRNA, and Prp28 have been studied in yeast via suppressor mutation analysis^13–15^. Kuhn *et al.* described a triple U4 snRNA mutation (*U4-cs1*) that stabilizes U4/U6 pairing and causes a cold-sensitive block in spliceosome activation, which can be rescued by *Prp8* mutations^16^. Similarly, *Prp8* variants can rescue the *brr2-1* phenotype, in which a mutation in the Brr2 helicase blocks U4/U6 RNA duplex unwinding^17^. Prp8 also regulates Prp28, which mediates U1 snRNP release and U6 pairing with the 5′SS (5’SS)^18^. The cold-sensitive *prp28-1* mutant (G279E) disrupts splicing, but its defects are suppressed by Prp8 mutations in the N-terminal domain^18^.

**Figure 1:**
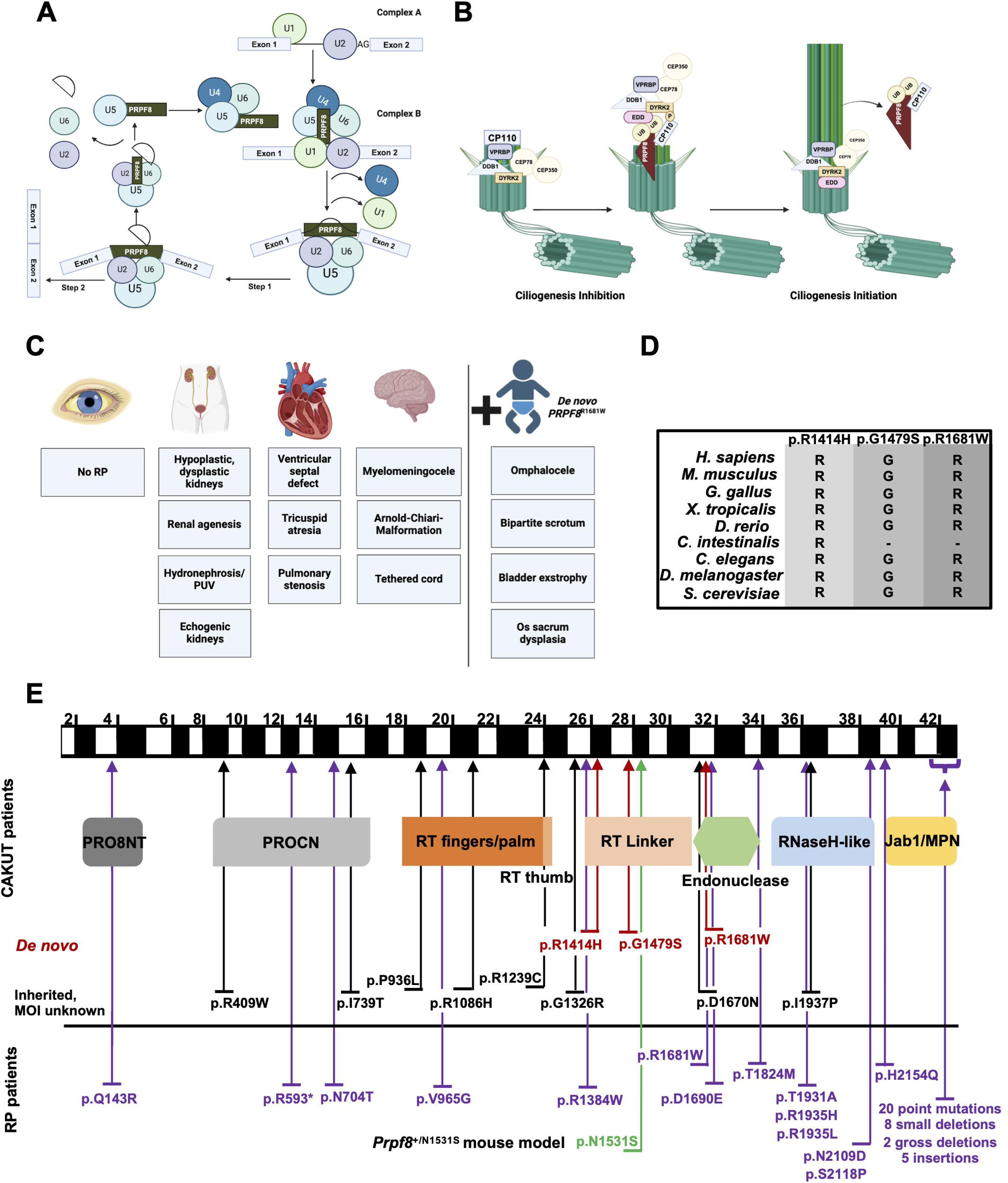
Schematic overview of PRPF8’s roles in splicing and ciliopathies. **A)** The graph provides an overview of the splicing process. PRPF8 is a core component of the U5 snRNP and the spliceosome. **B)** Ubiquitination and degradation of CP110 regulates ciliogenesis. CP110 caps distal end of the mother centriole and thereby inhibits ciliogenesis. EDD-DYRK2-DDB1^VprBP^ complex is constitutively located at the (sub)distal end of the mother centriole. CEP350 recruits CEP78, and CEP78 activates VPRBP and the EDD-DYRK2-DDB1^VprBP^ complex. Phosphorylation of CP110 by DYRK2 enables recognition of CP110, which is brought close to EDD that transfers ubiquitin to CP110. PRPF8 functions as receptor for ubiquitin chains of CP110. Ubiquitination causes CP110-disassembly and removal from mother centriole, initiating ciliogenesis. **C)** Summary of renal and extra-renal manifestations in patients with *PRPF8* variants. The patient with the *de novo PRPF8*^R1681W^ variant displayed the most severe phenotype with multiple malformations. None of the patients presented with RP symptoms. **D)** Examples of sequence conservation of PRPF8 amino acids mutated in CAKUT. **E)** Protein domain structure of human PRPF8 showing the position of *de novo* variants (red), heterozygous CAKUT variants (black) and RP mutations (magenta). Green arrow depicts a missense mutation (*Prpf8*^+/N1531S^) in a mouse model exhibiting a ciliopathy phenotype. PRO8NT: PRP8 N-terminal domain or Bromodomain; PROCN PRO8 central domain; RT reverse transcriptase domain; RNaseH-like Ribonuclease H domain; Jab1/MPN Jun activation domain-binding protein 1/Mpr1, Pad1 N-terminal domain; RP Retinitis Pigmentosa.

Recent studies suggest PRPF8 also plays a role in ciliogenesis. An siRNA screen by Wheway *et al.* (2016) identified *PRPF8* as a ciliopathy candidate gene, localizing to both the nucleus and ciliary base in mouse and human cells^19^. In *C. elegans*, a homozygous splice site variant in the *PRPF8* orthologue caused defective ciliogenesis^19^. Separately, Boylan *et al.* reported a mouse model carrying a missense variant in *Prpf8* (*Prpf8*^N1531S/N1531S^), displaying severe ciliopathy features, including left–right axis defects, open neural tube, heart malformations, and embryonic lethality at E10.5 with kidney necrosis^20^. *Prpf8*^N1531S/N1531S^ mice showed decreased *Shh, Gli1* and *Gli2* expression levels^21^, indicating impaired Sonic hedgehog (Shh) signaling- a pathway dependent on cilia^22^. Supporting a cilia-related role, PRPF8 was recently implicated in CP110 removal from the distal end of the mother centriole^5^, a key step in initiating ciliogenesis **(Fig. 1B)**^23^. This process involves the EDD-DYRK2-DDB1^VprBP^ E3 ubiquitin ligase and the linear ubiquitin chain assembly complex (LUBAC)^5, 24,25^. The EDD-DYRK2-DDB1^VprBP^-complex was proposed to induce CP110 removal via DYRK2-mediated phosphorylation^25^ and CEP78-dependent recruitment of EDD **(Fig. 1B)**^24^. PRPF8 may assist CP110 removal by acting as a receptor for LUBAC-generated linear ubiquitin chains^5^. Interestingly, during splicing, Prp8 is transiently ubiquitinated, and blocking this interaction or removing ubiquitin accelerates U4/U6 duplex unwinding, suggesting a regulatory role for Prp8 ubiquitination in splicing^26^.

Given its pivotal role in pre-mRNA splicing, *PRPF8* has been linked to several human diseases, including myeloid malignancies, primary open-angle glaucoma, and retinitis pigmentosa (RP)^11,27,28^. Similarly, pathogenic *CEP78* variants have been identified in patients with atypical Usher syndrome and RP^29^. Ascari *et al.* reported a *CEP78*^L150S^ variant in three families with cone-rod dystrophy and hearing loss^30^. Goncalves *et al.* later showed that this variant abrogates CEP78’s interaction with the EDD-DYRK2-DDB1^VprBP^ complex, and its recruitment to the centrosome via CEP350^24^. Supporting a ciliopathy-related role for this complex, Yoshida *et al.* showed that *Dyrk2*-deficient mice display congenital anomalies, including renal hypoplasia^31,32^. Furthermore, *de novo DDB1* variants have been implicated in syndromic phenotypes involving CAKUT^33^. Despite these findings, the precise role of *PRPF8, DYRK2, DBB1, CEP78,* and *EDD* variants (EDD-DYRK2-DDB1^VprBP^ complex) in human ciliopathies remains elusive.

Here, we identified heterozygous, both inherited and *de novo,* variants in the above-mentioned genes in patients with kidney malformations, suggesting they may cause CAKUT via disrupted splicing and/or ciliogenesis. Yeast assays showed that while *PRPF8* variants do not affect splicing at consensus splice sites, the *de novo* variant *PRPF8*^R1681W^ impairs splicing of non-consensus sites. CAKUT variants did not rescue *prp28-1* or *U4-cs1* cold-sensitive alleles but showed synthetic sickness at higher temperatures and weak suppression of *brr2-1*. To assess the impact of *PRPF8* variants on ciliogenesis, we measured Shh signaling and found that ∼50% of the variants impaired pathway activity, as indicated by reduced *GLI1* and *PTCH1* expression in SAG-treated RPE-1 cells. Two *DYRK2* variants impaired EDD-DYRK2-DDB1^VprBP^ complex formation. In situ hybridization revealed *Prpf8* expression throughout the embryonic urinary tract, later restricted to the collecting duct and urothelium. Our findings suggest that variants in *PRPF8,* and the EDD-DYRK2-DDB1^VprBP^ complex contribute to CAKUT/ciliopathy-like phenotypes by disrupting splicing and/or ciliogenesis.

## METHODS

### Research subjects, exome sequencing and variant calling

This study was approved by IRBs at the University of Michigan and Boston Children’s Hospital. Following consent, DNA and clinical data were collected from patients with isolated or syndromic CAKUT and their parents. Exome sequencing was performed on blood or saliva samples as previously described^34^; using Agilent SureSelect and Illumina HiSeq. Variants were aligned to hg19, filtered for rarity and potential pathogenicity, and analyzed for *de novo*, homozygous, or compound heterozygous changes. To conduct a thorough variant evaluation, we adhered to a decision-making strategy outlined in a previously published protocol^23^. Variant interpretation followed established protocols, incorporating prediction tools, conservation, structure modeling (AlphaFold^35^ and Robetta^36^), and literature review.

### Yeast strains and site-directed mutagenesis

Yeast strains and plasmids used in this study are listed in **Tables S1** and **S2**. Yeast transformation, plasmid shuffling/5-FOA selection, and growth were carried out using standard procedures^37,38^. *PRP8* mutants were constructed by site-directed mutagenesis using inverse PCR and the resulting plasmids were fully sequenced^39^.

### ACT1-CUP1 copper tolerance assays

Yeast strains containing WT or mutant *PRP8* and expressing ACT1-CUP1 reporters were grown to stationary phase in -Leu DO media to maintain selection for plasmids. Overnight cultures were diluted to OD_600_ = 0.5 in 10% (v/v) sterile glycerol before being spotted onto - Leu DO plates containing 0 to 2.5 mM CuSO_4_^40,41^. Plates were scored and imaged after 48 h of growth at 30°C.

### Yeast growth assays

To study the impact of temperature on yeast strain growth, strains were grown overnight in YPD media at 30°C before being diluted and stamped onto YPD plates. Plates were typically incubated for 3 days (23, 30, or 37°C) or 10 days (16°C) before imaging. For growth assays in the presence of 5-FOA, the same procedure was used except that yeast were plated on synthetic media lacking tryptophan in the presence (-TRP +5-FOA; 1 g/L) or absence (-TRP) of 5-FOA.

### Plasmids and cloning procedures

WT and mutant plasmids for human *PRPF8, CEP78, EDD,* and *DDB1* were obtained from GenScript. WT DYRK2 was purchased from OriGene, and mutants were generated using the QuikChange II XL kit (Agilent). *DYRK2* constructs were cloned with an N-terminal FLAG tag. EGFP-*CEP78* and Myc-*CEP350*-N plasmids were described previously^24,42^. *CEP78* variant plasmids were generated by GenScript in a pcDNA3.1-N-DYK backbone, excised with *Bam*H1/*Kpn*1, and ligated into pEGFP-C1 (TaKaRa Bio, cat. #6084-1).

### Cell lines and mouse fibroblasts

Immortalized RPE-1 (CRL-4000) and HEK293T (CRL-3216) cells were obtained from ATCC (hTERT RPE-1, CRL-4000 ™, HEK 293T cat. # CRL-3216). RPE-1 cells were cultured in DMEM/F-12 with hygromycin B, 10% FBS, and antibiotics; Lenti-X 293T (Takara Bio) in DMEM with 10% FBS, L-glutamine, and antibiotics. Transient transfections were performed using PEI Max on collagen I-coated dishes, and lysates were collected after 24 h for IP or western blotting. All cells were maintained at 37 °C with 5% COL. Fibroblasts from WT and *Prpf8*^N1531S/+^ mice (gift from K. Hentges) were cultured in DMEM with 10% FBS and antibiotics^20^.

### Shh signaling assay

RPE-1 cells were transfected with CAKUT variants plasmids using Lipofectamine™ 2000. Mouse fibroblasts and RPE-1 cells were serum-starved in Opti-MEM (Thermo Fisher) for 24 hours, followed by smoothened agonist (SAG) treatment to induce Shh signaling. RNA was isolated (RNeasy Mini Kit, Qiagen), reverse-transcribed (Superscript III, Invitrogen) and analyzed by qPCR using Sybr Green (Qiagen) on a MyiQ system (Bio-Rad Laboratories, Inc). Expression was normalized to 18s and *PRPF8* to exclude artificial effects due to differences in expression levels.

### DYRK2 phosphorylation assay

DYRK2 kinase activity was assessed using an EGFP-NDEL1-over-expression system^43^. Lenti-X 293T cells overexpressing EGFP-NDEL1 were transfected with each N-FLAG-*DYRK2* mutant and WT and measured by immuno-blotting using phospho-NDEL1^S3^^36^ antibody. Measurements were normalized to GAPDH. For the generation of an NDEL1^S3^^36^ phosphorylation-specific antibody, the peptide NH2-Cys-SSRPS(pS)APGML-COOH (> 80% purity) was obtained from SCRUM Inc. and used to immunize rabbits. Phosphorylation-specific IgGs were subsequently purified and then subjected to absorption with a non-phosphorylated peptide (SCRUM Inc.). To serve as a positive control, we employed the DYRK2^K251R^ mutant, a variant known for its ability to abrogate kinase function. An empty plasmid served as a negative control.

### Co-IP experiments of *DYRK2* and *CEP78* variants

To assess DYRK2 variant effects on EDD-DYRK2-DDB1VprBP complex formation, FLAG-tagged DYRK2 constructs were overexpressed in Lenti-X 293T cells. Lysates were prepared in NP-40 buffer with inhibitors, and immunoprecipitation was performed using FLAG M2 beads (Sigma-Aldrich). After washing and elution, proteins were analyzed by SDS-PAGE and western blotting. Signals were detected using ECL reagents and quantified via Fusion-Solo (M&S Instruments). For co-IP of EGFP-CEP78 variants and Myc-CEP350-N, HEK293T cells were co-transfected and lysed as previously described^24^. GFP-Trap (ChromoTek) was used for IP. Input/pellet fractions were immunoblotted using anti-Myc (CST), anti-VPRBP (Bethyl), and anti-GFP (Sigma) antibodies. Band intensities were quantified in Fiji from three replicates^44^.

### Protein complex structure prediction by AlphaFold multimer

For predicting protein complexes containing CEP78, CEP350 and DYRK2 we have used a local installation of AlphaFold multimer^45,46^. The CEP78-DYRK2 complex was folded from their full-length amino acid sequences. Because of its large size, the CEP350 amino acid sequence was initially split into its N- and C-terminal regions to facilitate computing and allow the prediction of complexes with CEP78. Ultimately, short fragments of CEP350 were used to predict complexes with CEP78 in different stoichiometries as depicted in **Fig. 5**.

### RNA in situ hybridization

A *Prpf8* cDNA subcloned into a pBluescript II KS+ plasmid was obtained from GensScript (Piscataway, NJ, USA). The cDNA had a length of 2,998 bps and represents the sequence between position 4,326 and 7,323 (the 3’-end of the open reading frame) of the mouse *Prpf8* mRNA (NCBI reference sequence: NM_138659.2). The plasmid was linearized with the restriction enzyme *XbaI* (BioLabs, Ipswich, MA, USA). For antisense RNA probe synthesis T3 RNA polymerase and the DIG-RNA labeling mix (Roche, Basel, Switzerland) were used. RNA in situ hybridization was performed on 10 µm paraffin-embedded sections of Naval Medical Research Institute (NMRI) mouse embryo trunks following a published protocol^47^. Sections were photographed using a Leica DM5000 microscope with Leica DFC300FX digital camera. Figures were assembled with Adobe Photoshop CS4.

### Statistical analysis

Statistical analysis was performed using GraphPad Prism 10.1.0. Data are shown as mean□±□SD. One-way ANOVA with Dunnett’s multiple comparisons test was used; significance: *P□≤□0.05, **P□≤□0.01, ***P□≤□0.001, ****P□≤□0.0001.

## RESULTS

### *PRPF8* variants in children with CAKUT

Trio exome sequencing in 208 CAKUT families revealed a *de novo PRPF8* variant. Literature review identified two additional CAKUT cases with *de novo PRPF8* variants^48,49^ **(Suppl. Table 1)**. Screening our broader cohort uncovered 8 more heterozygous *PRPF8* missense variants -either inherited or of unknown inheritance. Of 11 total cases, 8 showed isolated CAKUT (e.g., dysplastic kidneys, agenesis, hydronephrosis), while 3 had extra-renal features including cardiac and neurological malformations (**Fig. 1C, Suppl. Table 1**). The *de novo PRPF8*^R1681W^ variant was associated with the most severe phenotype, including omphalocele, bladder exstrophy, sacral dysplasia, tethered cord, and Arnold-Chiari malformation **(Fig. 1C, Suppl. Table 1**). *PRPF8* shows 61% sequence conservation with yeast *Prp8*, and all CAKUT variants affect highly conserved residues **(Fig. 1D)**^11^.

Interestingly, CAKUT variants cluster in exons 1–37, while 80.5% of RP variants are in exons 39–43 encoding the RNaseH-like and JAB1/MPN domains **(Fig. 1E)**. All *de novo* and 3 of 8 additional CAKUT variants map to the linker or endonuclease domains. **(Fig. 1E)**.

### CAKUT-associated *Prp8* mutants are viable in yeast

PRPF8 is a core spliceosome component. We hypothesized that CAKUT *PRPF8* variants impair splicing. Using yeast growth and ACT1-CUP1 splicing assays, we found that all variants supported growth at 16□°C and 30□°C but showed temperature-sensitive defects at 37□°C. The *de novo* variant *PRPF8*^R1681W^ (*Prp8*^R1753W^) was inviable at 37 °C (**Fig. 2A**).

**Fig. 2:**
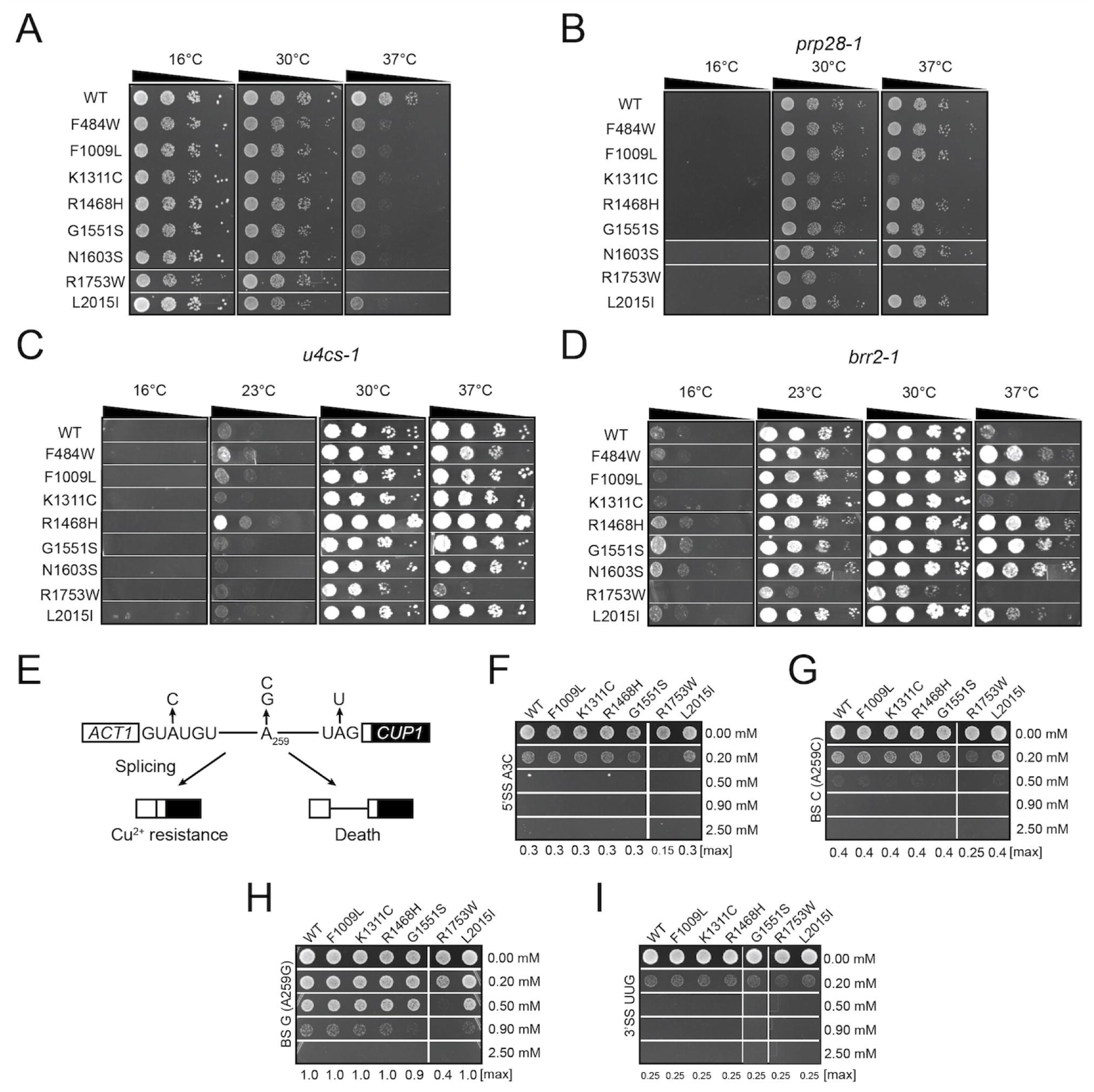
Splicing effects of *PRP8* variants. **A)** Growth of yeast containing different *Prp8* variants at various temperatures. A strain harboring the *de novo* variant *PRPF8*^R1681W^ (*Prp8*^R1753W^) is inviable at 37 °C. **B)** CAKUT variants do not rescue the *prp28-1* cold sensitive phenotype at 16 °C. **C)** Prp8 variants do not suppress the *U4-cs1* phenotype at 16 °C, however *PRPF8^R1414H^ (Prp8^R1468H^)* shows a weak suppression of *U4-cs1* at 23°C. **D)** Multiple *Prp8* variants show genetic interactions with *brr2-1*. **E)** Schematic overview of the ACT-1CUP1 assay, including the locations of non-consensus substitutions. **F-H)** ACT1-CUP1 assay results for the A3C (panel F), BS-C (panel G), BS-G (panel H), and UUG reporters (panel I).

### *PRPF8* CAKUT variants perturb Prp28 activity during transfer of 5’ splice site base pairing

The cold-sensitive *prp28-1* mutant (G279E) impairs splicing and spliceosome assembly. While known *Prp8* suppressors can rescue this phenotype, we tested whether CAKUT-associated *PRPF8* variants do the same^50^. None restored growth at 16□°C, indicating no suppression of prp28-1 (**Fig. 2A**, first column). However, *PRPF8*^K1239C^ (*Prp8*^K1311C^) and *PRPF8*^R1681W^ (*Prp8*^R1753W^) were synthetically sick with *prp28-1* at 30 °C and 37 °C suggesting that these mutants may perturb Prp28 activity during transfer of 5’ splice site base pairing from the U1 snRNA to the U6 snRNA (**Fig. 2B**, second/third column).

### *Prp8* CAKUT mutants may help to promote U4 snRNA release from the spliceosome

Removal of U4 snRNA is a prerequisite for formation of the spliceosome active site and achieved by disruption of the U4/U6 snRNA duplex by the Brr2 helicase. The cold sensitive mutant, *U4-cs1*, harbors a triple nucleotide substitution in the U4 snRNA and inhibits U4/U6 unwinding and spliceosome active site assembly at low temperatures (16 °C)^14^. Next, we tested if *Prp8* CAKUT variants are *U4-cs1* suppressors. *PRPF8^R1414H^ (Prp8^R1468H^)* showed suppression at 23 °C, while none of the tested variants suppressed the *U4-cs1* phenotype at 16 °C **(Fig. 2C)**. This suggests that some *Prp8* CAKUT mutants may help to promote U4 snRNA release from the spliceosome at 23 °C, while not being able to efficiently suppress defects that occur at lower temperatures.

### Multiple *Prp8* CAKUT-associated variants may affect spliceosome activation via Prp28- and Brr2-dependent processes

Brr2 promotes spliceosome activation by unwinding U4 from U6 snRNA, a process regulated by Prp8’s C-terminal domains (Rnase H-like and Jab1/MPN-like domains). The *brr2-1* mutation disrupts this activity by impairing the helicase domain, blocking U4/U6 duplex unwinding^13^. We tested whether *Prp8* CAKUT variants rescued the *brr2-1* phenotype. *PRPF8^R1414H^ (Prp8^R1468H^), PRPF8^G1479S^ (Prp8^G1551S^)* and *PRPF8^N1531S^ (Prp8^N1603S^)* were weak suppressors at 16 °C and strong suppressors at 37 °C **(Fig. 3D)**. These observations are not true for all suppressors or under all conditions. For example, the *PRPF8^12^*^39^ *(Prp8^K1311C^)* mutant is synthetically sick with *brr2-1* at 37 °C **(Fig. 2D)**. Together with our data from *prp28-1* and *U4-cs1* suppressors this suggests that CAKUT mutants can impact assembly of the spliceosome active site, potentially by altering Prp28-dependent splice site transfer and/or Brr2-dependent U4/U6 snRNA unwinding.

**Fig. 3:**
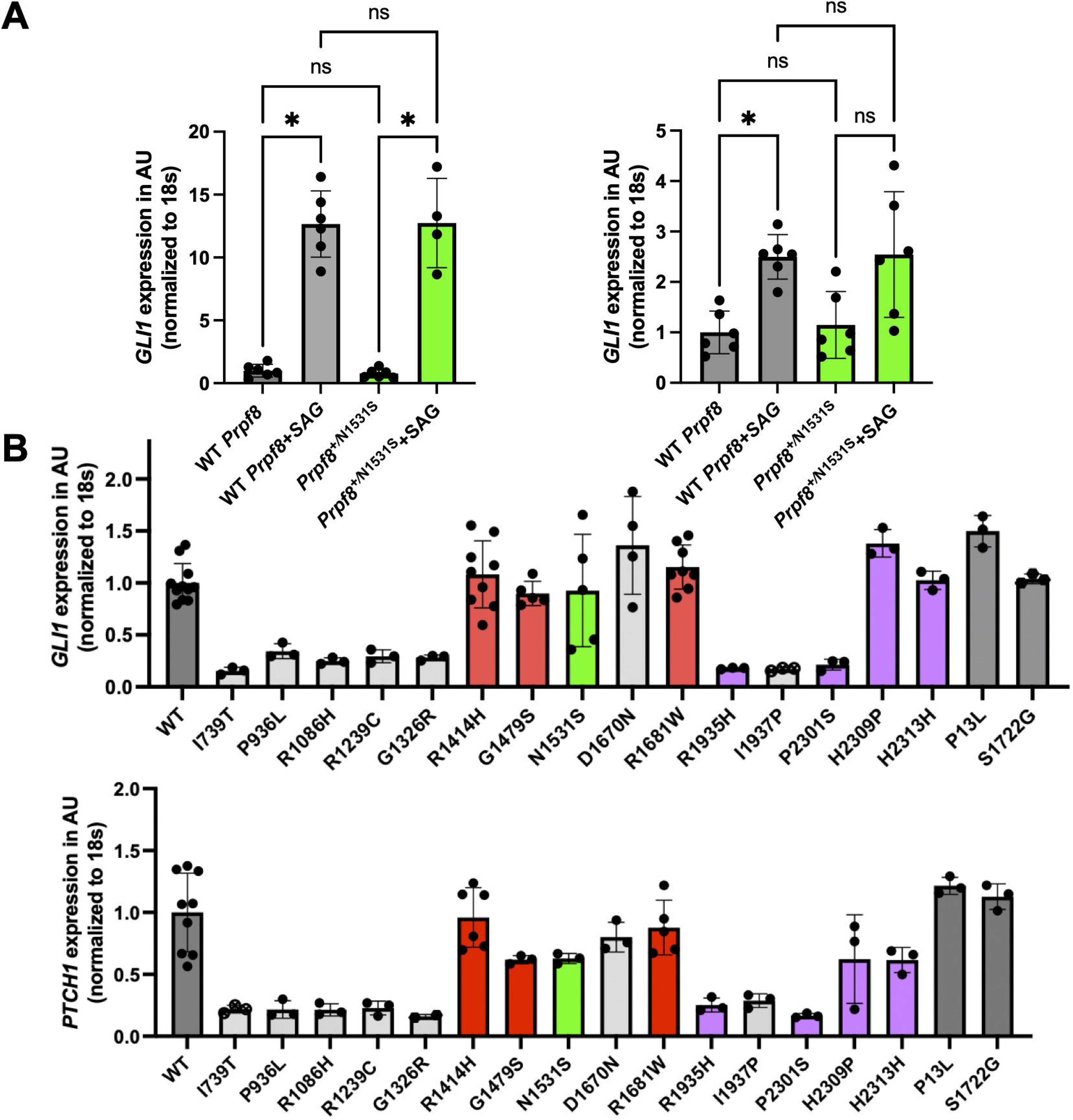
Effects of *PRPF8* variants on Shh signaling. **A)** No difference between ability to increase *Gli1* and *Ptch* expression in *Prpf8^+/N1531S^* mouse embryonic fibroblast treated with SAG. **B)** *GLI1* and *PTCH1* expression of RPE-1 cells transfected with plasmids endoding CAKUT variants (**red**=*de novo*, **green**=*Prpf8^N1531S^*, **grey**=CAKUT heterozygous), and RP variants **(magenta)**. Cells were stimulated with SAG and *GLI1* and *PTCH1* expression analyzed by RT-qPCR, using endogenous *PRPF8* mRNA and 18S RNA levels for normalization. P13L and S1722G serve as negative controls.

### *PRPF8*^R1681W^ (*Prp8*^R1753W^) shows reduced splicing activity

To assess splicing activity of *Prp8* CAKUT variants, we used a ACT1-CUP1 yeast reporter assay, where copper tolerance reflects correct pre-mRNA splicing **(Fig. 2E)**^40^. All variants showed normal splicing with consensus splice sites (**Suppl. Fig. 1**), indicating maintained splicing activity for this substrate in the presence of strong splice sites. Next, we used reporters with non-consensus sites affecting specific splicing steps^51^. With the BS-C reporter (A259C), *PRPF8*^R1681W^ (*Prp8*^R1753W^) showed reduced copper tolerance, indicating impaired 5′SS cleavage **(Fig. 2G)**. With the BS-G variant (A259G), tolerance was further reduced, suggesting an additional defect in exon ligation **(Fig. 2H)**. Finally, we tested A3C and UUG substitutions at the 5’ and 3’ SS, respectively **(Fig. 2F, 2I)**. The A3C 5′SS reporter that is limiting for exon ligation showed reduced tolerance, while the UUG 3′SS reporter did not (**Fig. 2I**), indicating that the CAKUT variants are not impacted by non-consensus 3’ SS usage. However, as with the other reporters, *PRPF8*^R1681W^ (*Prp8*^R1753W^) showed decreased copper tolerance with the A3C reporter (**Fig. 2F**). Together, these results suggest that *PRPF8*^R1681W^ (*Prp8*^R1753W^)] impairs both splicing steps at non-consensus splice sites.

### Effects of *PRPF8* variants on Shh signalling

Boylan *et al.* reported reduced Shh signaling in *Prpf8*^+/N1531S^ mice^20^. We, hence, tested Shh pathway activity in *Prpf8*^+/N1531S^ fibroblasts but found no difference in *Gli1* and *Ptch1* expression compared to WT after SAG stimulation **(Fig. 3A)**. We then expressed WT, CAKUT, or RP *PRPF8* variants in RPE-1 cells and measured SAG-induced *GLI1/PTCH1* expression. Six of 11 CAKUT variants-located in reverse transcriptase (fingers/palm, thumb), RNaseH-like, and Jab1/MPN domains-showed reduced expression. Variants in the reverse transcriptase linker and endonuclease domains, including all *de novo* variants, had no effect **(Fig. 3B)**.

### *EDD, DBB1* and *DYRK2* as potential CAKUT/ciliopathy candidate genes

Exome analysis of 228 CAKUT patients revealed *de novo* variants in *EDD* and *DDB1*. The EDD-DYRK2-DDB1^VprBP^-complex promotes ciliogenesis by phosphorylation and ubiquitylation of CP110 at the distal end of the mother centriole^24,25^. PRPF8 interacts with the EDD-DYRK2-DDB1^VprBP^ complex and functions as a receptor for LUBAC-generated linear ubiquitin chains on CP110, aiding its removal **(Fig. 1B)**^5^. CEP78 interacts with VPRBP, and is required for EDD-DYRK2-DDB1 complex recruitment to the mother centriole **(Fig. 1B)**^24,25^. Re-analysis of unsolved CAKUT and ciliopathy cases, Gene Matcher queries, and literature research, uncovered additional variants in *EDD, DBB1, DYRK2* and *CEP78* **(Suppl. Table 2-5)**.

Trio analysis identified a *de novo DDB1* variant in a patient with left kidney agenesis and right vesicoureteric reflux grade III, without extra-renal manifestations. White *et al.* published three CAKUT families with *de novo* variants in *DDB1*, exhibiting intellectual disability, facial dysmorphism, obesity, syndactyly, and multiple additional malformations **(Suppl. Table 3**)^33^. We found six additional heterozygous *DDB1* variants in isolated CAKUT cases. Similarly, six patients carried heterozygous *DYRK2* missense variants **(Suppl. Table 4)**. Four patients harbored *CEP78* variants, presenting with CAKUT and/or ciliopathy-like phenotypes-predominantly including dysplastic kidneys **(Suppl. Table 5)**. One patient (B2496, c.960C>G, p.320*) exhibited a typical syndromic ciliopathy phenotype with CAKUT, obesity, RP, and polydactyly. However, upon interrogating sequencing data for variants in established ciliopathy genes, we additionally identified a homozygous pathogenic *BBS12* variant (B2496, c.8139_8140dup, p.Phe2714Valfs*16), indicating that *CEP78* likely was not the sole cause for the clinical phenotype observed in this family. In contrast, in the second family (F752, c.1372G>T, p.458*) that presented with bilateral dysplastic kidneys, no pathogenic variant in any known ciliopathy gene was found **(Suppl. Table 5).**

### Alpha-fold predicts a direct interaction between CEP350 and CEP78

We identified 4 heterozygous variants in *CEP78:* two missense (c.283C>T, p.R95C; c.284G>A, p.R95H) and two nonsense variants (c.960C>G, p.Tyr320*, c.1372G>T, p.Glu458*). Using AlphaFold, we first aimed to predict interaction sites between CEP78 and CEP350. With a high prediction confidence, AlphaFold showed that CEP78 forms a homo-dimer interacting with two copies of CEP350. This interaction is predicted to be mediated by residues 810-850 in CEP350 and residues 518-565 in CEP78, respectively **(Fig. 4A-B)**.

**Fig. 4:**
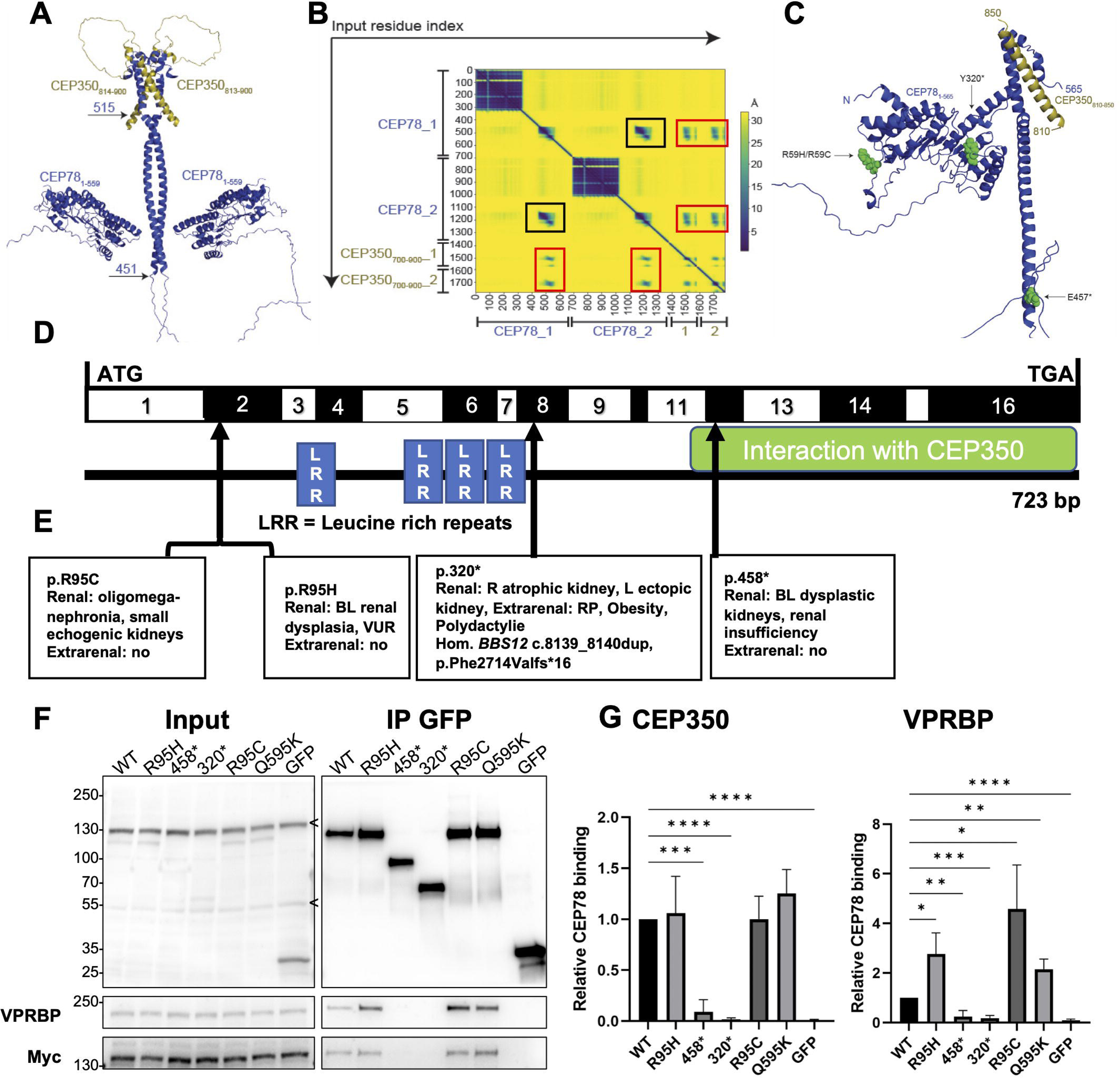
CEP350 is predicted to directly interact with CEP78 in a hetero-tetramer. **A)** AlphaFold predicts that a CEP78 dimer interacts with two CEP350 molecules via short helices (CEP350: residues 810–850; CEP78: residues 518–565). **B)** Truncating variants (p.320*, p.458*) eliminate the CEP350-binding region. **C)** Predicted aligned error (PAE) plot showing high-confidence interactions: black squares for CEP78 dimerization, red for CEP78– CEP350 binding. **D)** Exon structure of human *CEP78* cDNA. Arrowheads show variant position of CAKUT patients. **E)** Phenotypic features of four patients carrying a heterozygous *CEP78* variant. **F)** Co-IP of GFP-CEP78 variants and Myc-CEP350-N in HEK293T cells. Truncating variants show reduced size and no detectable interaction with VPRBP or CEP350. **G)** Band intensity ratios (VPRBP and Myc-CEP350 over GFP-IP) normalized to WT. Truncating variants abolish both interactions; missense variants retain CEP350 binding, with R95C and R95H showing increased VPRBP interaction.

### Truncating variants in *CEP78* abolish binding to VPRBP and CEP350

Based on the AlphaFold prediction, we hypothesized that truncating variants in *CEP78* completely abolish its interaction with CEP350 **(Fig. 4C)**. Hossain *et al.* recently described that CEP78 directly associates with VPRBP^25^. Moreover, Gonçalves *et al.* showed that CEP78 binds, at least indirectly, to the N-terminal region of CEP350 (residues 1–983) and to VPRBP^24^. We hypothesized that variants in *CEP78* may contribute to kidney abnormalities by affecting its interaction with CEP350 and VPRBP. To test our hypothesis, we co-expressed GFP-*CEP78* and Myc-*CEP350*-N in HEK293T cells and performed GFP IP analyses, followed by western blotting (*n=*3). Here, both truncating variants completely abolished binding capacity of CEP78 to CEP350-N and to VPRBP **(Fig. 4C-G)**. The *CEP78* missense variants, while not altering interaction with CEP350, did show significantly increased binding to VPRBP in comparison to WT *CEP78* **(Fig. 4C-G).**

### Two variants in *DYRK2* reduce EDD-DYRK2-DDB1^VprBP^-complex formation

Using AlphaFold we modeled DYRK2 interactions within the EDD-DYRK2-DDB1^VprBP^ complex and assessed whether CAKUT variants cluster at interaction sites. While a direct interaction with CEP78 was predicted, variants were dispersed throughout the protein and did not cluster **(Fig. 5 A-B)**. DYRK2 has been proposed to phosphorylate CP110, promoting its association to EDD. Four of six variants lie within *DYRK2*’s kinase domain **(Fig. 5A)**. We hypothesized that these variants might impair kinase function and used an NDEL1-based phosphorylation assay^43^. We overexpressed *DYRK2* variants in Lenti-X 293T cells and phospho-NDEL1^S3^^36^ concentrations were compared to WT. Interestingly, all variants retained phosphorylation capacity **(Fig. 5C)**.

**Fig. 5:**
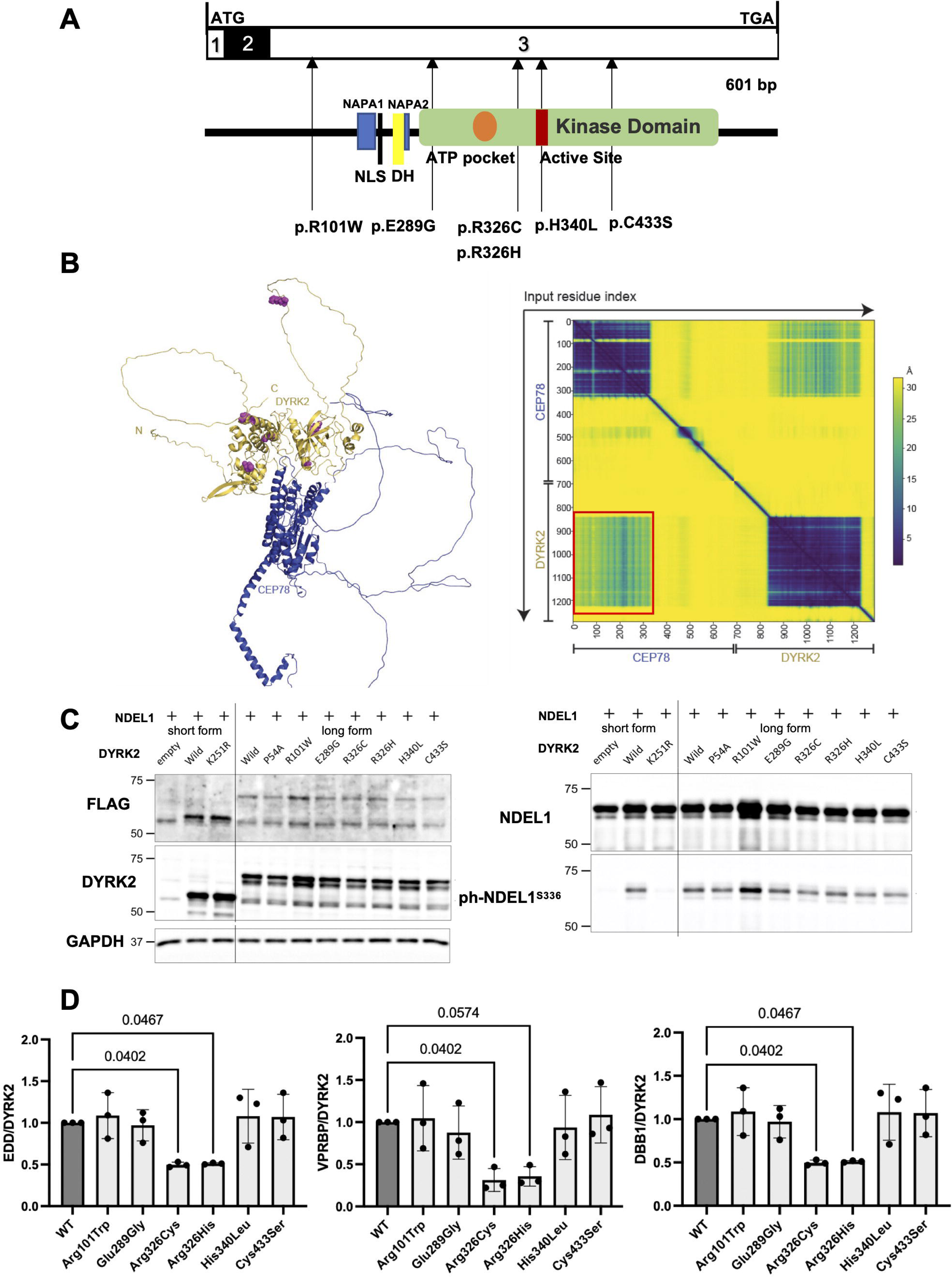
Two *DYRK2* CAKUT variants show decreased EDD-DYRK2-DDB1^VprBP^ complex formation efficiency. **A)** Exon and domain structure of human *DYRK2*. Arrowheads show positions of heterozygous CAKUT variants. **B)** Alphafold-predicted 3D structure of the DYRK2–CEP78 complex. The red square in the PAE plot suggests a potential direct interaction. Variants are scattered and do not cluster at the interface. **C)** DYRK2 kinase activity assessed via NDEL1 phosphorylation. Immunoblotting shows comparable phospho-NDEL1S336 levels for all CAKUT variants relative to WT, indicating preserved kinase function. K251R served as a kinase-dead control; empty vector as negative control. **D)** Co-IP of *DYRK2* variants with EDD, DDB1, and VprBP. Two variants (p.Arg326Cys, p.Arg326His) showed reduced complex formation (n=3).

Maddika *et al.* showed that *DYRK2* mediates EDD-DYRK2-DDB1^VprBP^ complex formation independent of its kinase activity^52^. We postulated that *DYRK2* variants disrupt this assembly. Co-IP of overexpressed variants revealed that two missense variants near the kinase domain’s ATP pocket (p.Arg326Cys, p.Arg326His) reduced complex formation (n=3, **Fig. 5D)**.

### *Prpf8* is strongly expressed in the developing murine urinary system

Having identified *PRPF8* variants in subjects with CAKUT/ciliopathy-like phenotypes, we performed *in situ* RNA hybridization in mouse to investigate *Prpf8* expression during kidney development. Interestingly, *Prpf8* is strongly and ubiquitously expressed until E14.5, in the entire urinary tract system **(Fig. 6)**. Expression declines at E16.5 and E18.5; however, expression persists in the collecting duct epithelium and the urothelium **(Fig. 6)**. These results are consistent with an important role for PRPF8 in kidney development and function.

**Fig. 6:**
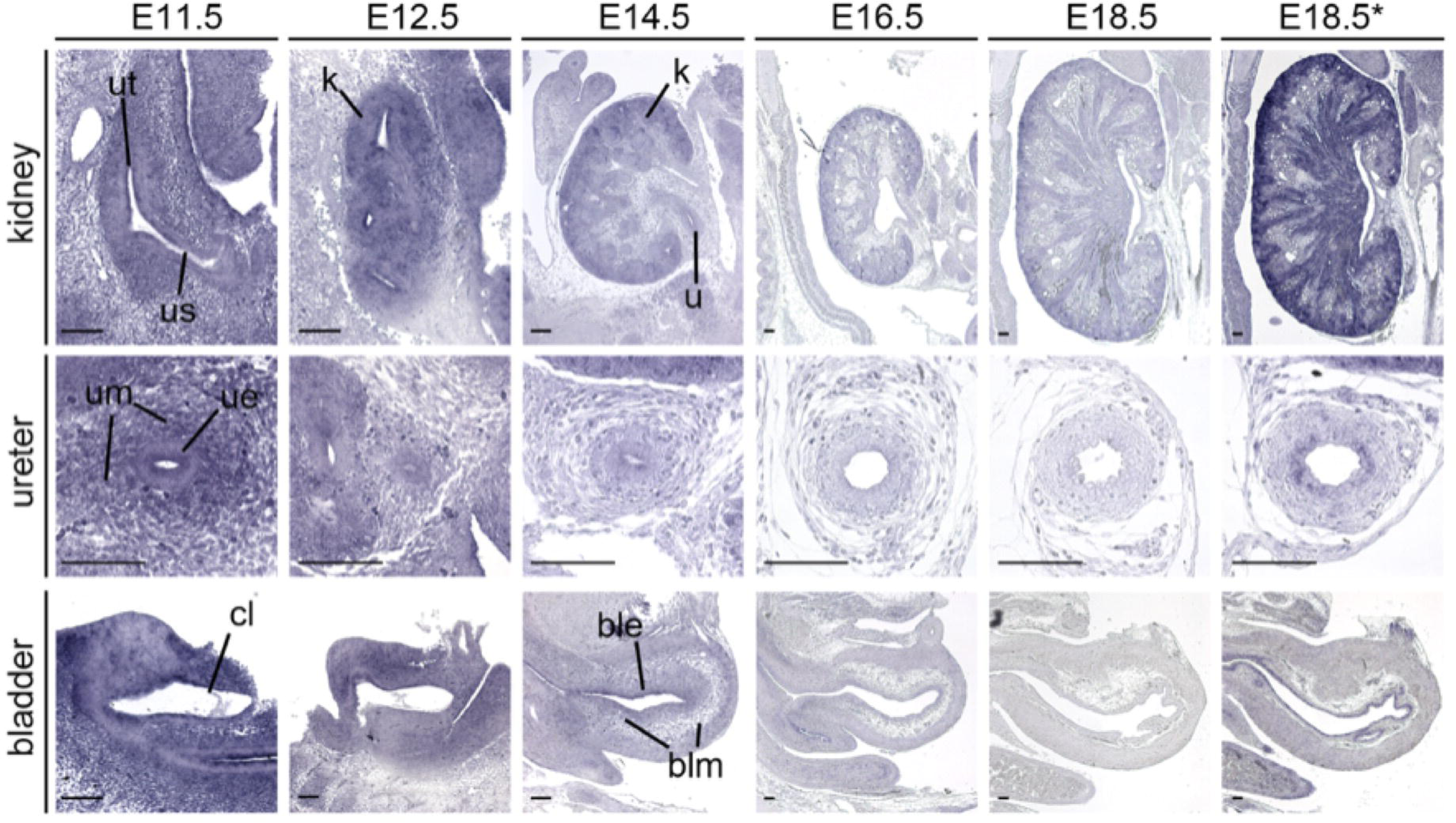
P*r*pf8 is widely expressed in the urinary system until E14.5. RNA *in situ* hybridization analysis of *Prpf8* expression on sagittal sections of mouse kidney (first row), transverse sections of the ureter (second row) and sagittal sections of the bladder (third row) of wildtype embryos from E11.5 to E18.5. Note that all sections were developed for the same time, except the ones from the last column (*) for which color development was prolonged to detect weak expression domains. n*=4* for each stage and tissue. Size bars represent 100 µm. ble, bladder epithelium; blm, bladder mesenchyme; cl, cloaca; k, kidney; u, ureter; ue, ureteric epithelium; um, ureteric mesenchyme; us, ureteric stalk, ut, ureteric tip.

## DISCUSSION

In this study, trio exome sequencing in CAKUT patients identified heterozygous variants in *PRPF8* and interactors at the distal mother centriole. Yeast splicing assays showed the *de novo* variant *PRPF8*^R1681W^ (*PRP8*^R1753W^) significantly reduces splicing capacity with different non-consensus splice sites. CAKUT variants did not rescue *prp28-1* or *U4-cs1* cold sensitive phenotypes, yet some of the variants showed synthetic sickness at higher temperatures and weakly suppressed *brr2-1*. As an indirect measure of ciliary dysfunction, Shh signaling was assessed in SAG-treated RPE-1 cells expressing *PRPF8* variants, and approximately 50% showed reduced *GLI1* and *PTCH1* expression. Two *DYRK2* variants disrupted formation of the EDD–DYRK2–DDB1VprBP complex. In situ hybridization revealed *Prpf8* expression throughout the urinary tract until E14.5, later becoming restricted to the collecting duct epithelium and urothelium.

PRPF8 shows strong missense constraint (Z-score□=□11.34, gnomAD), indicating high intolerance to variation, supporting autosomal dominant inheritance and pathogenicity of novel missense variants. Yeast expressing the CAKUT-*PRPF8* (*Prp8)* variants remain viable, suggesting these variants do not cause complete loss of function in humans. However, homozygous *Prpf8*^N1531S/N1531S^ mice are embryonically lethal^20^. The absence of homozygous *PRPF8* variants in our cohort and in population databases, suggests such variants may also be lethal in humans.

PRPF8 is a highly conserved protein (98% identity between human and zebrafish) primarily known for its role in pre-mRNA splicing^53^. Heterozygous *PRPF8* variants have been linked to splicing defects in Retinitis pigmentosa (RP) and myeloid malignancies, however no kidney phenotype had been reported prior to this study^11,54^. Supporting its role in CAKUT, our *in situ* hybridization revealed strong *Prpf8* expression in the mouse collecting duct epithelium during later stages of development. Of note, disruption of this epithelium is known to cause hypo- or dysplastic kidneys, the predominant phenotype of the here described patients^55–57^.

RP is a genetically heterogeneous disease, causing progressive vision loss due to photoreceptor and/or retinal pigment epithelium degeneration, typically beginning in early adulthood. The vast majority of RP *PRPF8* variants cluster within the last exons, affecting the Jab1/MPN domain^58^. In contrast, *PRPF8* variants detected in CAKUT patients are distributed outside this domain, primarily within the reverse transcriptase (fingers, thumb, linker) and endonuclease regions. Hereby, phenotypic limitation for the eye of RP variants could be explained by either dominant-negative effects, gain of function or haploinsufficiency^59^. The distinct clustering of CAKUT versus RP variants suggests allelism and domain-specific effects, leading to different genotype-phenotype correlations. Supporting this, reverse phenotyping of all CAKUT individuals with *PRPF8* variants revealed no RP symptoms. However, since RP often manifests in early adulthood, continued follow-up is warranted.

Unlike other CAKUT variants, yeast expressing *Prp8*^R1753W^ is inviable at 37 °C, suggesting a stronger splicing defect. Interestingly, the individual harboring the *PRPF8*^R1681W^ variant (*Prp8*^R1753W^) presented with a severe multisystem phenotype, including abdominal, bone, cardiac and CNS malformations. Yeast cells expressing *Prp8*^R1753W^ exhibit copper tolerances similar to those with WT *Prp8*, when using reporters with consensus splice sites. In contrast, they show reduced copper tolerance with non-consensus 5’SS or branch sites-similar to the previously reported *PRP8*^R1753K^ allele-suggesting disruption of catalytic splicing steps^51^. However, as *PRP8*^R1753K^ is temperature sensitive, additional effects beyond these steps cannot be excluded. Some variants also alter spliceosome active site assembly, a process highly regulated by Prp8^60^. Several CAKUT variants interact with *prp28-1*, *u4-cs1*, and *brr2-1*, all of which affect this process, indicating that *PRPF8* variants may change splicing at numerous stages and through diverse mechanisms.

PRPF8 has recently been proposed to function as a receptor for ubiquitinated CP110 at the distal mother centriole, promoting its removal and initiating ciliogenesis^5^. EDD and DDB1, as components of the EDD-DYRK2-DDB1^VprBP^ ubiquitin E3 ligase complex, collaborate with PRPF8 in this process. Ciliopathies represent a genetically diverse disease group, ranging from single organ involvement to severe systemic disease^61^. Typical kidney phenotypes include hypoplastic, dysplastic, and echogenic kidneys, as well as CAKUT features such as kidney agenesis and horseshoe kidneys^62^. We hypothesized that CAKUT variants impair Shh signaling, however only approximately half of the *PRPF8* variants showed reduced pathway activity. A caveat with this assay is that endogenous WT *PRPF8* is still expressed in these cells at levels that may be sufficient for normal ciliogenesis and Shh signaling capacity.

Loss-of-function variants in *CEP78* cause a specific type of cone-rod dystrophy accompanied by hearing loss^63,64^. Over decades the mechanism of ciliogenesis regulation by CEP78 remained elusive. Recently, Gonçalves *et al.* showed that *CEP78* loss reduces ciliation frequency and increases residual cilia length^24^. They further demonstrated that CEP78 is recruited to the centrosome via CEP350 binding, promoting activation of the EDD-DYRK2-DDB1^VprBP^ complex. We detected heterozygous *CEP78* frameshift variants in patients with kidney ciliopathy/CAKUT phenotype, which completely abrogated CEP78 binding to CEP350 and VPRBP. As these variants (p.320*, p.458*) truncate the CEP350-binding region (residues 518–565), even if they escape nonsense-mediated decay, the resulting proteins are likely non-functional. Also, reverse phenotyping revealed no symptoms of Usher syndrome, encompassing vision and hearing loss. Re-analysis of a syndromic ciliopathy case (B2496, p.320*) harboring a *CEP78* variant revealed a homozygous pathogenic *BBS12* variant (**Fig. 4E, Suppl. Table 5**). Bardet-Biedl syndrome (BBS) is genetically heterogeneous, with a significant level of inter- and intra-familial variability. While typically considered an autosomal recessive disease with variants in one of the 24 BBS genes, oligogenic inheritance patterns are increasingly discussed for BBS pathogenesis and might in part explain the observed heterogeneity. Triallelism was first reported two decades ago^65^, and a recent study suspected an oligogenic inheritance in 52% of 45 screened BBS families^66^. In such cases, BBS gene variants act as primary drivers, with additional variants modifying disease severity^67,68^. *CEP78*, involved in cilia formation, interacts with CEP350 and the EDD-DYRK2-DDB1^VprBP^ complex to promote CP110 degradation. Truncating *CEP78* variants may impair this process, causing CP110 accumulation and defective ciliogenesis, potentially contributing to or modifying a ciliopathy phenotype.

In summary, our data suggests that variants in *PRPF8* and the EDD-DYRK2-DDB1^VprBP^ complex can cause a kidney ciliopathy/CAKUT phenotype in humans. While recent studies highlight their interaction, the role of PRPF8 in ciliogenesis remains incompletely understood. Splicing assays indicate that *PRPF8* variants disrupt splicing at multiple stages, suggesting that kidney phenotypes may result from defective pre-mRNA splicing and/or impaired ciliogenesis.

## ACKNOWLEDGEMENTS

We thank all participating families, physicians, and collaborators for their invaluable contributions.

## CONFLICT OF INTEREST

AAH is a member of the SAB for Remix Therapeutics and is carrying out sponsored research in collaboration with Remix. All other authors declare no conflicts of interest.

## DATA SHARING STATEMENT

The data that support the findings of this study are available from the corresponding author upon reasonable request. Due to ethical and privacy considerations, sequencing data and detailed patient information are not deposited in a public repository at this stage. All relevant functional and experimental data are included within the manuscript and supplementary files. Additional materials may be shared for academic, non-commercial purposes upon reasonable request, following institutional and ethical guidelines.

## FUNDING STATEMENT

F. H. is the William E. Harmon Professor of Pediatrics at Harvard Medical School. This research was supported by grants from the National Institutes of Health (NIH) to F.H. (DK076683) and by the Begg Family Foundation. L.M.M. (456136540) and C.M.K. (499462148, Rückkehrstipendium KO 6579/ /3-1) are supported by the German Research Foundation (Deutsche Forschungsgemeinschaft, DFG). N.D.M was supported by the NIH (5T32-DK007726-37) and the Fred Lovejoy House-staff Research and Education Fund. F.B. is supported by the Else Kröner-Fresenius-Stiftung (iPRIME Clinician Scientist Forschungskolleg - 2021_EKFK.15, UKE, Hamburg, Germany) and received funding through the Carl W. Gottschalk Research Scholar Grant from the American Society of Nephrology. L.B.P acknowledges funding from the Novo Nordisk Foundation (grant NNF18SA0032928 and NNF22OC0080406), Independent Research Fund Denmark (grant 2032-00115B), and the European Union’s Horizon 2020 research and innovation program Marie Sklodowska-Curie Innovative Training Networks (ITN) grant 861329. A.A.H. was supported by funding from the National Institutes of Health (R35 GM136261) with additional support from a Research Forward grant award from the Wisconsin Alumni Research Foundation. A.C.H. was supported by the Else Kröner-Fresenius Stiftung and the Eva Luise und Horst Köhler Stiftung (Project No: 2019_KollegSE.04). Work in the laboratory of A.K. is supported by grants from the German Research Foundation (DFG KI728/10-2, DFG KI728/12-1). M.A.C. and R.J.J. were supported by Grant PID2021-124314OB-I00, funded by MICIU/AEI/10.13039/501100011033 and co-funded, as applicable, by ‘ERDF – A Way of Making Europe,’ ‘ERDF/EU,’ the ‘European Union,’ or the ‘European Union NextGenerationEU/PRTR’ program. This research was supported by British Heart Foundation grants PG/06/144/21898, PG/10/87/28624 and PG/18/28/33632 to K.E.H.

## ETHICS DECLARATION

The institutional review boards at both the University of Michigan and Boston Children’s Hospital, along with those at the recruiting institutions, approved this study. Written informed consent was acquired from each participant or their legal guardians prior to inclusion in the study.

**Fig. S1: *Prp8* variants do not change splicing at consensus splice sites. A)** Schematic representation of plasmid transformation in yeast. **B)** CAKUT variants are viable in yeast. **C)** *Prp8* variants and WT exhibit similar copper tolerances when the 5’splice site consensus reporter is used.

## Supplementary Material

Supplementary Table 1-9

Supplementary Figure 1

## LITERATURE

1. Van Der Ven AT, Vivante A, Hildebrandt F. Novel insights into the pathogenesis of monogenic congenital anomalies of the kidney and urinary tract. J Am Soc Nephrol. 2018;29(1):36–50. doi:10.1681/ASN.2017050561

2. Lossky M, Anderson GJ, Jackson SP, Beggs J. Identification of a yeast snRNP protein and detection of snRNP-snRNP interactions. Cell. 1987;51(6):1019–1026. doi:10.1016/0092-8674(87)90588-5

3. Stevens SW, Barta I, Ge HY, et al. Biochemical and genetic analyses of the U5, U6, and U4/U6.U5 sStevens, S. W., Barta, I., Ge, H. Y., Moore, R. E., Young, M. K., Lee, T. D., & Abelson, J. (2001). Biochemical and genetic analyses of the U5, U6, and U4/U6.U5 small nuclear ribonucleoproteins. Rna. 2001;7(11):1543–1553.

4. Gottschalk A, Neubauer G, Banroques J, Mann M, Lührmann R, Fabrizio P. Identification by mass spectrometry and functional analysis of novel proteins of the yeast [U4/U6·U5] tri-snRNP. EMBO J. 1999;18(16):4535–4548. doi:10.1093/emboj/18.16.4535

5. Shen XL, Yuan JF, Qin XH, et al. LUBAC regulates ciliogenesis by promoting CP110 removal from the mother centriole. J Cell Biol. 2022;221(1). doi:10.1083/jcb.202105092

6. Seltzsam S, Wang C, Zheng B, et al. Reverse phenotyping facilitates disease allele calling in exome sequencing of patients with CAKUT. Genet Med. 2022;24(2):307–318. doi:10.1016/j.gim.2021.09.010.Reverse

7. Vidal VPI, Verdone L, Mayes AE, Beggs JD. Characterization of U6 snRNA-protein interactions. Rna. 1999;5(11):1470–1481. doi:10.1017/S1355838299991355

8. Maroney PA, Romfo CM, Nilsen TW. Functional recognition of the 5’ splice site by U4/U6.U5 tri-snRNP defines a novel ATP-dependent step in early spliceosome assembly. Mol Cell. 2000;6(2):317–328. doi:10.1016/S1097-2765(00)00032-0

9. Dix I, Russell CS, O’Keefe RT, Newman AJ, Beggs JD. Protein-RNA interactions in the U5 snRNP of Saccharomyces cerevisiae. RNA. 1998;4(12):1675–1686.

10. McPheeters DS. Interactions of the yeast U6 RNA with the pre-mRNA branch site. Rna. 1996;2(11):1110–1123.

11. Kurtovic-Kozaric A, Przychodzen B, Singh J, et al. PRPF8 Defects Cause Missplicing in Myeloid Malignancies. Leukemia. 2015;29(1):126–136. doi:doi:10.1038/leu.2014.144.

12. López-Cánovas JL, Hermán-Sánchez N, del Rio-Moreno M, et al. PRPF8 increases the aggressiveness of hepatocellular carcinoma by regulating FAK/AKT pathway via fibronectin 1 splicing. Exp Mol Med. 2023;55(1):132–142. doi:10.1038/s12276-022-00917-7

13. Galej WP, Oubridge C, Newman AJ, Nagai K. Crystal structure of Prp8 reveals active site cavity of the spliceosome. Nature. 2013;493(7434):638-643. doi:10.1038/nature11843

14. Kuhn AN, Brow DA. Suppressors of a cold-sensitive mutation in yeast U4 RNA define five domains in the splicing factor Prp8 that influence spliceosome activation. Genetics. 2000;155(4):1667–1682. doi:10.1093/genetics/155.4.1667

15. Kuhn AN, Reichl EM, Brow DA. Distinct domains of splicing factor Prp8 mediate different aspects of spliceosome activation. Proc Natl Acad Sci U S A. 2002;99(14):9145–9149. doi:10.1073/pnas.102304299

16. Kuhn AN, Li Z, Brow DA. Splicing factor Prp8 governs U4/U6 RNA unwinding during activation of the spliceosome. Mol Cell. 1999;3(1):65–75. doi:10.1016/S1097-2765(00)80175-6

17. Nielsen KH, Staley JP. Spliceosome activation: U4 is the path, stem I is the goal, and Prp8 is the keeper. let’s cheer for the ATPase Brr2! Genes Dev. 2012;26(22):2461–2467. doi:10.1101/gad.207514.112

18. Yeh FL, Chang SL, Ahmed GR, et al. Activation of Prp28 ATPase by phosphorylated Npl3 at a critical step of spliceosome remodeling. Nat Commun. 2021;12(1):1–9. doi:10.1038/s41467-021-23459-4

19. Wheway G, Schmidts M, Mans DA, et al. An siRNA-based functional genomics screen for the identification of regulators of ciliogenesis and ciliopathy genes□: Nature Cell Biology□: Nature Publishing Group. NatureCom. 2016;17(8):1074–1087. doi:10.1038/ncb3201.An

20. Boylan M. A Novel Point Mutation in Prpf8 Causes Defects in Left-Right Axis Establishment in the Mouse 2015. Univ Manchester. Published online 2015.

21. Stephen LA. Identification and characterisation of cardiac defects in mouse models isolated from a random chemical mutagenesis screen. Univ Manchester. Published online 2013.

22. Bangs F, Anderson K V. Primary cilia and Mammalian Hedgehog signaling. Cold Spring Harb Perspect Biol. 2017;9(5):1–21. doi:10.1101/cshperspect.a028175

23. Xie S, Naslavsky N, Caplan S. Emerging insights into CP110 removal during early steps of ciliogenesis. J Cell Sci. 2024;137(4):jcs261579. 10.1242/jcs.261579

24. Gonçalves AB, Hasselbalch SK, Joensen BB, et al. CEP78 functions downstream of CEP350 to control biogenesis of primary cilia by negatively regulating CP110 levels. Elife. 2021;10:1–34. doi:10.7554/eLife.63731

25. Hossain D, Javadi Esfehani Y, Das A, Tsang WY. Cep78 controls centrosome homeostasis by inhibiting EDD □ DYRK 2□ DDB 1 Vpr BP. EMBO Rep. 2017;18(4):632–644. doi:10.15252/embr.201642377

26. Bellare P, Small EC, Huang X, Wohlschlegel JA, Staley JP, Sontheimer EJ. A role for ubiquitin in the spliceosome assembly pathway. Nat Struct Mol Biol. 2008;15(5):444–451. doi:10.1038/nsmb.1401

27. McKie AB, McHale JC, Keen TJ, et al. Mutations in the pre-mRNA splicing factor gene PRPC8 in autosomal dominant retinitis pigmentosa (RP13). Hum Mol Genet. 2001;10(15):1555–1562. doi:10.1093/hmg/10.15.1555

28. Micheal S, Hogewind BF, Khan MI, et al. Variants in the PRPF8 Gene are Associated with Glaucoma. Mol Neurobiol. 2018;55(5):4504–4510. doi:10.1007/s12035-017-0673-5

29. L L, S H, S T, et al. A novel frameshift variant in CEP78 associated with nonsyndromic retinitis pigmentosa, and a review of CEP78-related phenotypes. Ophthalmic Genet. 2022;43(2):152–158. doi:10.1080/13816810.2022.2045511.

30. Ascari G, Peelman F, Farinelli P, et al. Functional characterization of the first missense variant in CEP78, a founder allele associated with cone-rod dystrophy, hearing loss, and reduced male fertility. Hum Mutat. 2020;41(5):998–1011. doi:10.1002/humu.23993

31. Yoshida S, Aoki K, Fujiwara K, et al. The novel ciliogenesis regulator dyrk2 governs hedgehog signaling during mouse embryogenesis. Elife. 2020;9:1–29. doi:10.7554/ELIFE.57381

32. Yoshida S, Kawamuraa A, Aokib K, et al. Positive regulation of Hedgehog signaling via phosphorylation of GLI2/GLI3 by DYRK2 kinase. Proc Natl Acad Sci U S A. 2024;121(28). doi:doi: 10.1073/pnas.2320070121. Epub 2024 Jul 5.

33. White SM, Bhoj E, Nellåker C, et al. A DNA repair disorder caused by de novo monoallelic DDB1 variants is associated with a neurodevelopmental syndrome. Am J Hum Genet. 2021;108(4):749–756. doi:10.1016/j.ajhg.2021.03.007

34. Braun DA, Sadowski CE, Kohl S, et al. Mutations in nuclear pore genes NUP93, NUP205 and XPO5 cause steroid-resistant nephrotic syndrome. Nat Genet. 2016;48(4):457–465. doi:10.1038/ng.3512

35. AlphaFold Protein Structure Database. https://alphafold.ebi.ac.uk

36. Robetta. https://robetta.bakerlab.org

37. Treco DA, Lundblad V. Preparation of Yeast Media. Curr Protoc Mol Biol. 1993;23(1):1–7. doi:10.1002/0471142727.mb1301s23

38. Sikorski RS, Boeke JD. In vitro mutagenesis and plasmid shuffling: From cloned gene to mutant yeast. Methods Enzymol. 1991;194(1989):302-318.

39. Green MR, Sambrook J. Inverse polymerase chain reaction (PCR). Cold Spring Harb Protoc. 2019;2019(2):170–174. doi:10.1101/pdb.prot095166

40. Lesser CF, Guthrie C. Mutational analysis of pre-mRNA splicing in Saccharomyces cerevisiae using a sensitive new reporter gene, CUP1. Genetics. 1993;133(4):851–863. doi:10.1093/genetics/133.4.851

41. Carrocci TJ, Paulson JC, Hoskins AA. Functional analysis of Hsh155/SF3b1 interactions with the U2 snRNA/branch site duplex. Rna. 2018;24(8):1028–1040. doi:10.1261/rna.065664.118

42. Eguether T, Ermolaeva MA, Zhao Y, et al. The deubiquitinating enzyme CYLD controls apical docking of basal bodies in ciliated epithelial cells. Nat Commun. 2014;5(May). doi:10.1038/ncomms5585

43. Woo Y, Kim SJ, Suh BK, et al. Sequential phosphorylation of NDEL1 by the DYRK2-GSK3β complex is critical for neuronal morphogenesis. Elife. 2019;8:1–23. doi:10.7554/eLife.50850

44. Schindelin J, Arganda-Carreras I, Frise E, et al. Fiji: an open-source platform for biological-image analysis. Nat Methods. 2012;9:676–682. doi:10.1038/nmeth.2019

45. Jumper J, Evans R, Pritzel A, et al. Highly accurate protein structure prediction with AlphaFold. Nature. 2021;596(7873):583–589. doi:10.1038/s41586-021-03819-2

46. Evans R, O’Neill M, Alexander Pritzel, et al. Protein complex prediction with AlphaFold-Multimer. bioRxiv Prepr. Published online 2022. 10.1101/2021.10.04.463034

47. Moorman AFM, Houweling AC, De Boer PAJ, Christoffels VM. Sensitive nonradioactive detection of mRNA in tissue sections: Novel application of the whole-mount in situ hybridization protocol. J Histochem Cytochem. 2001;49(1):1–8. doi:10.1177/002215540104900101

48. Lei TY, Fu F, Li R, et al. Whole-Exome Sequencing in the Evaluation of Fetal Congenital Anomalies of the Kidney and Urinary Tract Detected by Ultrasonography. Vol 40.; 2020. doi:10.1002/pd.5737

49. Reutter H, Keppler-Noreuil K, E. Keegan C, Thiele H, Yamada G, Ludwig M. Genetics of Bladder-Exstrophy-Epispadias Complex (BEEC): Systematic Elucidation of Mendelian and Multifactorial Phenotypes. Curr Genomics. 2015;17(1):4–13. doi:10.2174/1389202916666151014221806

50. Grainger RJ, Beggs JD. Prp8 protein: At the heart of the spliceosome. Rna. 2005;11(5):533–557. doi:10.1261/rna.2220705

51. Liu L, Query CC, Konarska MM. Opposing classes of prp8 alleles modulate the transition between the catalytic steps of pre-mRNA splicing. Nat Struct Mol Biol. 2007;14(6):519–526. doi:10.1038/nsmb1240

52. Maddika S, Chen J. Protein kinase DYRK2 is an E3-ligase specific molecular assembler. Nat Cell Biol. 2009;11(4):409–419. doi:10.1038/ncb1848.Protein

53. Luo HR, Moreau GA, Levin N, Moore MJ. The human Prp8 protein is a component of both U2- and U12-dependent spliceosomes. Rna. 1999;5(7):893–908. doi:10.1017/S1355838299990520

54. Mayerle M, Guthrie C. Prp8 retinitis pigmentosa mutants cause defects in the transition between the catalytic steps of splicing. Rna. 2016;22(5):793–809. doi:10.1261/rna.055459.115

55. Walton SL, Singh RR, Little MH, Bowles J, Li J, Moritz KM. Prolonged prenatal hypoxia selectively disrupts collecting duct patterning and postnatal function in male mouse offspring. J Physiol. 2018;596(23):5873–5889. doi:10.1113/JP275918

56. Piscione TD, Rosenblum ND. The malformed kidney: disruption of glomerular and tubular development. Clin Genet. 1999;56(5):341–356. doi:10.1034/j.1399-0004.1999.560502.x.

57. Kozlov VM, Schedl A. Duplex kidney formation: Developmental mechanisms and genetic predisposition. F1000Research. 2020;9:1–12. doi:10.12688/f1000research.19826.1

58. Wang J, Xiao X, Li S, et al. Landscape of pathogenic variants in six pre-mRNA processing factor genes for retinitis pigmentosa based on large in-house data sets and database comparisons. Acta Ophthalmol. 2022;100(7):e1412–e1425. doi:10.1111/aos.15104

59. Daniel Mordes, Yuan L, Xu L, Kawada M, Molday RS, Wu JY. Identification of photoreceptor genes affected by PRPF31 mutations associated with autosomal dominant retinitis pigmentosa. Neurobiol Dis. 2007;26(2):291–300.

60. Brow DA. Allosteric cascade of spliceosome activation. Annu Rev Genet. 2002;36:333–360. doi:10.1146/annurev.genet.36.043002.091635

61. Focşa IO, Budişteanu M, Bălgrădean M. Clinical and genetic heterogeneity of primary ciliopathies. Int J Mol Med. 2021;48(3):1–15. doi:10.3892/ijmm.2021.5009

62. Jenkins D, PL Beales. Genes and mechanisms in human ciliopathies. In: Emery and Rimoin’s Principles and Practice of Medical Genetics. Academic Press, Oxford. ; 2013:pp1□36.

63. Q F, M X, X C, et al. CEP78 is mutated in a distinct type of Usher syndrome. J Med Genet. 2017;54(3):190–195. doi:10.1136/jmedgenet-2016-104166.CEP78

64. Namburi P, Ratnapriya R, Khateb S, et al. Bi-allelic Truncating Mutations in CEP78, Encoding Centrosomal Protein 78, Cause Cone-Rod Degeneration with Sensorineural Hearing Loss. Am J Hum Genet. 2016;99(3):777–784. doi:10.1016/j.ajhg.2016.07.010

65. Katsanis N, Ansley SJ, Badano JL, et al. Triallelic inheritance in Bardet-Biedl syndrome, a Mendelian recessive disorder. Science (80-). 2001;293(5538):2256–2259. doi:10.1126/science.1063525

66. Perea-Romero I, Solarat C, Blanco-Kelly F, et al. Allelic overload and its clinical modifier effect in Bardet-Biedl syndrome. npj Genomic Med. 2022;7(1):1–7. doi:10.1038/s41525-022-00311-2

67. Deltas C. Digenic inheritance and genetic modifiers. Clin Genet. 2018;93(3):429–438. doi:10.1111/cge.13150

68. Manara E, Paolacci S, D’esposito F, et al. Mutation profile of BBS genes in patients with Bardet-Biedl syndrome: An Italian study. Ital J Pediatr. 2019;45(1):1–8. doi:10.1186/s13052-019-0659-1

