## Supplementary figures and images for "CAKUT variants in *PRPF8, DYRK2*, and *CEP78*: implications for splicing and ciliogenesis"

### Supplementary Figure 1

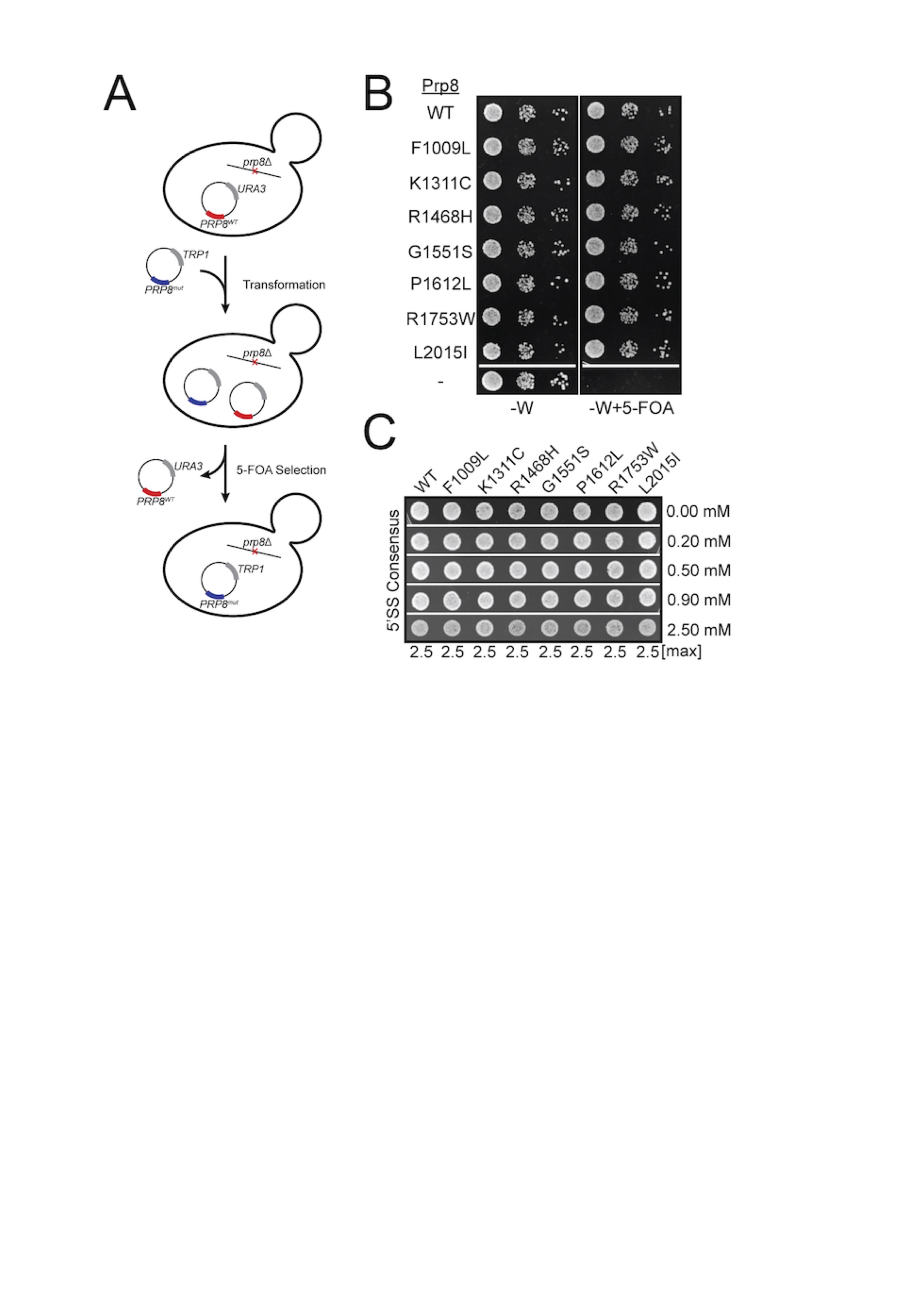
