## Supplementary Table 1 for "CAKUT variants in *PRPF8, DYRK2*, and *CEP78*: implications for splicing and ciliogenesis"

**Table 1. *De novo* and heterozygous mutations in *PRPF8* in 11 individuals with CAKUT/ciliopathy.**

| Family Number | Nucleotide/ amino acid change^a, b^ | Amino acid  conservation | PP2  SIFT  MUT | Zygosity | gnomAD allele frequency^a^ | Ethnicity  Gender | CAKUT | Extra-renal manifestations |
| --- | --- | --- | --- | --- | --- | --- | --- | --- |
| B4047 | c.4241G>A p.Arg1414His | S.c. | 0.95  D  DC | *de novo* | NP | Caucasian  F | Renal agenesis, unilateral | NP |
| B3185 | c.1225C>T p.Arg409Trp | D.m. | 0.92  T  DC | het | 0/1/31366 | Caucasian  F | BL UPJO, BL hydronephrosis | NP |
| B4225 | c.2806T>C p.Phe936Leu | S.c. | 0.99  D  DC | het | NP | Caucasian  F | Small lower pole of the right kidney, neurogenic bladder | Myelomeningocele, Arnold Chiari type 2 |
| B4025 | c.3715C>T  p.Arg1239Cys | D.m. | 0.98  D  DC | het | NP | Caucasian  F | Renal agenesis, unilateral, ectopic insertion of ureter into lower bladder neck | VACTERL syndrome: tricuspid atresia, subvalvular pulmonic stenosis, tethered cord |
| Collaborators | c.2216T>C p.Ile739Thr | D.m. | 1  D  DC | het | 0/1/251366 | Latino  M | Hypodysplastic kidneys | NP |
| Collaborators | c.3257G>A p.Arg1086His | S.c. | 1  D  DC | het | 0/2/251442 | Caucasian  M | Renal hypoplasia, unilateral | NP |
| Collaborators | c.3976G>C p.Gly1326Arg | D.m. | 1  D  DC | het | NP | Caucasian  M | UPJO, unilateral | NP |
| Collaborators | c.5008G>A p.Asp1670Asn | D.m. | 0.99  D  DC | het | NP | Caucasian  F | Renal hypoplasia, bilateral  VUR | NP |
| Collaborators | c.5809A>T p.Ile1937Phe | D.m. | 0.99  D  DC | het | NP | Caucasian  M | Renal dysplasia, bilateral | NP |
| Reutter *et al.*^1^ | c.5041C>T p.Arg1681Trp | D.m. | 0.99  D  DC | *de novo* | NP | Caucasian  M | Bladder exstrophy | Omphalocele, tethered cord, Arnold Chiari, ventricular septal defect, permanent foramen ovale, os sacrum dysplasia, biparte scrotum, inguinal hernia, epilepsy |
| Lei *et al.*^2^ | c.4435G>A  p.Gly1479Ser | D.m. | 1  D  DC | *de novo* | NP | NA | Multicystic dysplastic  kidney, unilateral | NP |

**Table 2. *De novo* and heterozygous variants in *EDD* in 7 individuals with CAKUT/ciliopathy.**

| Family Number | Nucleotide/ amino acid change^,^ | Amino acid  conservation | PP2  SIFT  MUT | Zygosity | gnomAD allele frequency^a^ | Ethnicity  Gender | CAKUT | Extra-renal manifestations |
| --- | --- | --- | --- | --- | --- | --- | --- | --- |
| B3208 | c.6737G>A  p.Arg2246Gln | D.m. | 0.98  T  DC | *de novo* | NP | Caucasian  M | Ureterocele  Left hydronephrosis | NP |
| B3418 | c.6473T>A  p.Leu2158His | D.m. | 0.97  T  DC | het | NP | Arabic  M | PUV, Left VUR Hydronephrosis bilateral | NP |
| A4670 | c.2461C>T  p.Pro821Ser | D.m. | 0.96  T  DC | het | NP | Caucasian  F | Bilateral VUR II-IV° | NP |
| A3367 | c.8153T>C  p.Val2718Ala | D.m. | 0.99  D  DC | het | NP | Caucasian  M | Right Renal agenesis | Cryptorchidism, enuresis |
| A4850 | c.3103C>T  p.Pro1035Ser | D.m. | 0.99  D  DC | het | NP | Caucasian  F | VUR bilateral | NP |
| B437 | c.2438G>A  p.Arg813Gln | D.m. | 0.88  D  DC | het | 0/3/251398 | Caucasian  M | Right small kidney | NP |
| B3199 | c.2759G>A  p.Arg920Gln | D.m. | D B | het | 0/1/251290 | Caucasian  M | Hypodysplastic right kidney, right  VUR II° | NP |

**Table 3. *De novo* and heterozygous mutations in *DDB1* in 9 individuals with CAKUT/ciliopathy.**

| Family Number | Nucleotide/ amino acid change^a, b^ | Amino acid  conservation | PP2  SIFT  MUT | Zygosity | gnomAD allele frequency^a^ | Ethnicity  Gender | CAKUT | Extra-renal manifestations |
| --- | --- | --- | --- | --- | --- | --- | --- | --- |
| B3163 | c.1688A>T  p.Asp563Val | S.c. | 0.92  T  DC | *de novo* | NP | Caucasian  M | L renal agenesis,  R VUR III° | NP |
| B4045 | c.2260C>T  p.Pro754Ser | C.i. | 0.02  T  DC | het | 0/2/250418 | Caucasian  F | L VUR III°,  L reflux nephropathy | NP |
| B720 | c.1685C>T  p.Thr562Met | D.m. | 0.5  D  DC | het | 0/1/251366 | African-  American  M | PUV, megaureters,  R dysplastic kidney, R VUR V° | NP |
| Collaborators | c.3224C>G  p.Ser1075Cys | C.i. | 0.99  D  DC | het | NP | Caucasian  M | R UPJO | NP |
| Collaborators | c.656T>C  p.Val219Ala | S.c. | 0.9  D  DC | het | NP | Caucasian  F | R VUR | NP |
| Collaborators | c.176G>C  p.Gly59Ala | S.c. | 1  T  DC | het | NP | Caucasian  F | L renal agenesis | NP |
| White *et al.*^3^ | c.563G>A  p.Arg188Gln | D.m. | 0.96  T  B | *de novo* | NP | NA  F | L horseshoe kidney with VUR pelvic dilatation, megaureter | mild-moderate ID, facial dysmorphism, short toes, mild joint laxity |
| White *et al.*^3^ | c.637G>A  p.Glu213Lys | D.m. | 0.96  T  B | *de novo* | NP | NA  M | L mild hydronephrosis | moderate ID, facial dysmorphism, 2–3 toe partial syndactyly, gastro-esophageal reflux |
| White *et al.*^3^ | 562C>T  p.Arg188Trp | D.m. | 1  D  DC | *de novo* | NP | NA  F | horseshoe kidney | mild ID, facial dysmorphism, accessory band across left ventricle of heart,  anterior ectopic anus, dysgenesis of corpus callosum, 2–3 toe syndactyly,  brachydactyly, obesity |

**Table 4. Heterozygous mutations in *DYRK2* in 6 individuals with CAKUT/ciliopathy.**

| Family Number | Nucleotide/ amino acid change^a, b^ | Amino acid  conservation | PP2  SIFT  MUT | Zygosity | gnomAD allele frequency^a^ | Ethnicity  Gender | CAKUT | Extra-renal manifestations |
| --- | --- | --- | --- | --- | --- | --- | --- | --- |
| B1861 | c.866A>G  p.Glu289Gly | C.i. | 0.99  D  DC | het | NP | Caucasian  F | Left UPJO, Left hydronephrosis  Echogenic kidney, SRNS | NP |
| B3247 | c.301C>T  p.Arg101Trp | D.r. | 0.93  D  DC | het | 0/4/251204 | Caucasian  M | VUR V°, unilateral | NP |
| B1486 | c.1019A>T  p.His340Leu | D.r. | 0.29  D  DC | het | 0/1/251475 | Egyptian  M | Ectopic right kidney, VUR II°  Hypoplastic kidney unilateral, VUR IV-V°  Megaureters, enlarged ureter | NP |
| Collaborators | c.976C>T  p.Arg326Cys | C.e. | 1  D  DC | het | 0/3/251377 | Caucasian  M | Renal hypoplasia, unilateral | NP |
| Collaborators | c.977G>A  p.Arg326His | C.e. | 0.97  D  DC | het | 0/1/251359 | Caucasian  M | Renal agenesis, unilateral  Hydronephrosis, unilateral | NP |
| Collaborators | c.1298G>C  p.Cys433Ser | D.m. | 1  D  DC | het | NP | Caucasian  M | Renal agenesis, unilateral  Hydronephrosis, unilateral, megaureter, VUR | NP |

**Table 5. Heterozygous mutations in CEP78 in 4 individuals with CAKUT/ciliopathy.**

| Family Number | Nucleotide/ amino acid change^a, b^ | Amino acid  conservation | PP2  SIFT  MUT | Zygosity | gnomAD allele frequency^a^ | Ethnicity  Gender | CAKUT | Extra-renal manifestations |
| --- | --- | --- | --- | --- | --- | --- | --- | --- |
| B1603 | c.283C>T  p.R95C | C.i. | 0.93  D  DC | het | 0/15/1581106 | Hispanic  M | Oligomeganephronia  small echogenic kidneys, bilateral | NP |
| B722 | c.284G>A  p.R95H | D.r. | 0.89  D  DC | het | 0/14/1581696 | Hispanic  M | Bilateral renal dysplasia, bilateral high-grade  VUR | NP |
| B2496  *BBS12*  *CEP78* | c.8139_8140dup  p.Phe2714Valfs*16  c.960C>G  p.320* | NA  NA | NA  NA | hom  het | NA  0/2/212278 | Arabic  M | Atrophic kidney, unilateral  Ectopic kidney, unilateral | RP, Obesity, Polydactyly |
| F752 | c.1372G>T  p.458* | NA | NA | het | NP | Caucasian  F | Hypodysplastic kidneys, bilateral  Renal insufficiency | NP |

**Supplemental Table S6.** Yeast strains used in this study.

| **Name** | **Genotype** | **Description** | **Additional**  **Variants** | **Reference** |
| --- | --- | --- | --- | --- |
| yAAH0117 (yJU75) | MATa, cup1Δ ura3 his3 leu2 lys2 prp8Δ:lys2 trp1 pJU169 (PRP8 URA3) | Cu^2+^-sensitive shuffle strain for Prp8 | -- | ^4^ |
| yAAH3093 | yAAH0117 +  (PRP8 TRP1 2µ) | Prp8^WT^ | Prp8-(F484W, F1009L, K1311C, R1468H, G1551S, N1603S, P1612L, R1753W, L2015I) on TRP1 CEN plasmids | ^5^ |
| yAAH3106 | yAAH2435+  pAAH0470 (LEU2) | Prp8^WT^ Cu^2+^-sensitive strain with WT ACT-CUP1 | Prp8-(F484W, F1009L, K1311C, R1468H, G1551S, N1603S, P1612L, R1753W, L2015I) on TRP1 CEN plasmids | ^5^ |
| yAAH3107 | yAAH2435+  pAAH1032 (LEU2) | Prp8^WT^ Cu^2+^-sensitive strain with A3C ACT-CUP1 | Prp8-(F484W, F1009L, K1311C, R1468H, G1551S, N1603S, P1612L, R1753W, L2015I) on TRP1 CEN plasmids | ^5^ |
| yAH3108 | yAAH2435+  pAAH0441 (LEU2) | Prp8^WT^ Cu^2+^-sensitive strain with A259C (BS-C) ACT-CUP1 | Prp8-(F484W, F1009L, K1311C, R1468H, G1551S, N1603S, P1612L, R1753W, L2015I) on TRP1 CEN plasmids | ^5^ |
| yAH3109 | yAAH2435+  pAAH0880 (LEU2) | Prp8^WT^ Cu^2+^-sensitive strain with A259G (BS-G) ACT-CUP1 | Prp8-(F484W, F1009L, K1311C, R1468H, G1551S, N1603S, P1612L, R1753W, L2015I) on TRP1 CEN plasmids | ^5^ |
| yAAH3110 | yAAH2435+  pAAH0527 (LEU2) | Prp8^WT^ Cu^2+^-sensitive strain with A302U (UUG) ACT-CUP1 | Prp8-(F484W, F1009L, K1311C, R1468H, G1551S, N1603S, P1612L, R1753W, L2015I) on TRP1 CEN plasmids | ^5^ |
| yAAH3156 (ANK828) | MATa prp28-1 prp8Δ::ADE2 trp1 ura3 his3 ade2 lys2 (YCp50-PRP8) | Genomic *prp28-1* strain with shuffle for Prp8 mutants | Prp8-(WT, F484W, F1009L, K1311C, R1468H, G1551S, N1603S, P1612L, R1753W, L2015I) on TRP1 CEN plasmids | ^6^ |
| yAAH3196 (ZRL102) | MATa snr14::TRP1::trp1Δ63 prp8Δ::ADE2 trp1 ura3 lys2 his3 ade2 [pRS317-U4-cs1] [yCp50-PRP8] | *u4cs-1* strain (on plasmid) with shuffle for Prp8 mutants | Prp8-(WT, F484W, F1009L, K1311C, R1468H, G1551S, N1603S, P1612L, R1753W, L2015I) on TRP1 CEN plasmids | ^7^ |
| yAAH3157 (ANK821) | MATa brr2-1 prp8Δ::ADE2 ura3 his3 ade2 lys2 (YCp50-PRP8) | Genomic *brr2-1* strain with shuffle for Prp8 mutants | Prp8-(WT, F484W, F1009L, K1311C, R1468H, G1551S, N1603S, P1612L, R1753W, L2015I) on TRP1 CEN plasmids | ^6^ |

**Supplemental Table S7**. Plasmids used in this study.

| **Plasmid ID** | **Plasmid name** | **Description** | **Source or reference** |
| --- | --- | --- | --- |
| pAAH1440 | Prp8^WT^ | Plasmid used to generate Prp8^WT^ and mutant strains (pRS424 PRP8 2micron TRP1) | Derived from pJU225-4^4^. |
| pAAH0470 | ACT1-CUP1 WT | WT reporter used for ACT1-CUP1 assays. ACT1-CUP1 expressed from a GPD promoter, LEU2 plasmid. | Gift from Charles Query. |
| pAAH1032 | ACT1-CUP1 A3C | 5′ss A3C reporter used for ACT1-CUP1 assays. ACT1-CUP1 expressed from a GPD promoter, LEU2 plasmid. | Gift from Charles Query. |
| pAAH0441 | ACT1-CUP1 BSC (A259C) | BS A259C reporter used for ACT1-CUP1 assays. ACT1-CUP1 expressed from a GPD promoter, LEU2 plasmid. | Gift from Charles Query. |
| pAAH0880 | ACT1-CUP1 BSG (A259G) | BS A259C reporter used for ACT1-CUP1 assays. ACT1-CUP1 expressed from a GPD promoter, LEU2 plasmid. | Gift from Charles Query. |
| pAAH0527 | ACT1-CUP1 A302U | 3′ss UuG reporter used for ACT1-CUP1 assays. ACT1-CUP1 expressed from a GPD promoter, LEU2 plasmid. | Gift from Charles Query. |
| pAAH1443 | Prp8^F484W^ | pRS424 PRP8-F484W 2micron TRP1 |  |
| pAAH 1444 | Prp8^F1009L^ | pRS424 PRP8-F1009L 2micron TRP1 |  |
| pAAH 1445 | Prp8^K1311C^ | pRS424 PRP8-K1311C 2micron TRP1 |  |
| pAAH 1446 | Prp8^R1486H^ | pRS424 PRP8-R1486H 2micron TRP1 |  |
| pAAH 1447 | Prp8^G1551S^ | pRS424 PRP8-G1551S 2micron TRP1 |  |
| pAAH 1448 | Prp8^P1612L^ | pRS424 PRP8-P1612L 2micron TRP1 |  |
| pAAH 1449 | Prp8^R1753W^ | pRS424 PRP8-R1753W 2micron TRP1 |  |
| pAAH 1450 | Prp8^L2015I^ | pRS424 PRP8-L2015I 2micron TRP1 |  |
| pAAH 1467 | Prp8^N1603S^ | pRS424 PRP8-N1603S 2micron TRP1 |  |
| U6755HE060 | CEP78^WT^  and Mutants | CEP78_OHu31091C_pcDNA3.1(+)-N-DYK | GenEZ ORF Clone from GenScript |
| OHu20334C | DDB1^WT^  and Mutants | DDB1_OHu20334C_pcDNA3.1(+)-N-HA | GenScript |
| U7598HG070 | EDD^WT^  and Mutants | UBR5_pcDNA3.1(+)-N-DYK | GenScript |
| OHu19527C | PRPF8^WT^  and Mutants | PRPF8_OHu19527C_pcDNA3.1(+)-N-Myc | GenScript |

**Supplemental Table S8.** Summary of Representative ACT1-CUP1 Results

|  | **ACT1-CUP1 Reporter, Maximum Cu^2+^ Tolerance (mM)** | | | | |
| --- | --- | --- | --- | --- | --- |
| **Prp8 Mutant** | **WT** | **5' SS A3C** | **BS C** | **BS G** | **3' SS UUG** |
| **WT** | 2.5 | 0.3 | 0.4 | 1.0 | 0.25 |
| **F484W** | 2.5 | 0.3 | 0.4 | 1.0 | 0.3 |
| **F1009L** | 2.5 | 0.3 | 0.4 | 1.0 | 0.25 |
| **K1311C** | 2.5 | 0.3 | 0.4 | 1.0 | 0.25 |
| **R1468H** | 2.5 | 0.3 | 0.4 | 1.0 | 0.25 |
| **G1551S** | 2.5 | 0.3 | 0.4 | 0.9 | 0.25 |
| **N1603S** | 2.5 | 0.3 | 0.25 | 0.4 | 0.3 |
| **P1612L** | 2.5 | 0.15 | 0.3 | 0.8 | 0.25 |
| **R1753W** | 2.5 | 0.15 | 0.25 | 0.4 | 0.25 |
| **L2015I** | 2.5 | 0.3 | 0.4 | 1.0 | 0.25 |

Differences >0.05 mM [Cu^2+^] from WT are indicated in red.

**Supplemental Table S9. Overview of applied antibodies.**

| **Antibody** | **Source** | **Identifier** |
| --- | --- | --- |
| Anti-DYRK2 (Rabbit polyclonal) | Sigma-Aldrich | Cat# HPA027230, RRID:AB_1847925 |
| Anti-GAPDH (Mouse monoclonal) | Santa Cruz Biotechnology | Cat# sc-32233, RRID:AB_627679 |
| Anti-DYKDDDDK (Mouse monoclonal) | Sigma-Aldrich | Cat# F3165, RRID:AB_259529 |
| Anti-NDEL1 (Rabbit polyclonal) | Proteintech | Cat# 17262-1-AP, RRID:AB_2235821 |
| Anti-phosNDEL1^S336^ (Rabbit polyclonal) | This paper | N/A |
| Anti-EDD1 (Mouse monoclonal) | Santa Cruz Biotechnology | Cat# sc-376860, RRID:AB_2894825 |
| Anti-VPRBP (Mouse monoclonal) | Santa Cruz Biotechnology | Cat# sc-376850, RRID:AB_2905506 |
| Anti-DDB1 (Mouse monoclonal) | Santa Cruz Biotechnology | Cat# sc-376860, RRID:AB_2894825 |

1.	Reutter H, Keppler-Noreuil K, E. Keegan C, Thiele H, Yamada G, Ludwig M. Genetics of Bladder-Exstrophy-Epispadias Complex (BEEC): Systematic Elucidation of Mendelian and Multifactorial Phenotypes. *Curr Genomics*. 2015;17(1):4-13. doi:10.2174/1389202916666151014221806

2.	Lei TY, Fu F, Li R, et al. *Whole-Exome Sequencing in the Evaluation of Fetal Congenital Anomalies of the Kidney and Urinary Tract Detected by Ultrasonography*. Vol 40.; 2020. doi:10.1002/pd.5737

3.	White SM, Bhoj E, Nellåker C, et al. A DNA repair disorder caused by de novo monoallelic DDB1 variants is associated with a neurodevelopmental syndrome. *Am J Hum Genet*. 2021;108(4):749-756. doi:10.1016/j.ajhg.2021.03.007

4.	Umen JG, Guthrie C. A novel role for a U5 snRNP protein in 3′ splice site selection. *Genes Dev*. 1995;9(7):855-868. doi:10.1101/gad.9.7.855

5.	Feltz C Van Der, Nikolai B, Schneider C, Paulson JC, Fu X, Hoskins AA. Saccharomyces cerevisiae Ecm2 modulates the catalytic steps of pre-mRNA splicing. *Rna*. 2021;27(5):591-603. doi:10.1261/rna.077727.120

6.	Kuhn AN, Reichl EM, Brow DA. Distinct domains of splicing factor Prp8 mediate different aspects of spliceosome activation. *Proc Natl Acad Sci U S A*. 2002;99(14):9145-9149. doi:10.1073/pnas.102304299

7.	Kuhn AN, Brow DA. Suppressors of a cold-sensitive mutation in yeast U4 RNA define five domains in the splicing factor Prp8 that influence spliceosome activation. *Genetics*. 2000;155(4):1667-1682. doi:10.1093/genetics/155.4.1667
